# *In vitro* and *in vivo* effects of zoledronate on senescence and senescence-associated secretory phenotype markers

**DOI:** 10.1101/2023.02.23.529777

**Authors:** Parinya Samakkarnthai, Dominik Saul, Lei Zhang, Zaira Aversa, Madison L. Doolittle, Jad G. Sfeir, Japneet Kaur, Elizabeth J Atkinson, James R. Edwards, R. Graham G. Russell, Robert J. Pignolo, James L. Kirkland, Tamar Tchkonia, Laura J. Niedernhofer, David G. Monroe, Nathan K. LeBrasseur, Joshua N. Farr, Paul D. Robbins, Sundeep Khosla

## Abstract

In addition to reducing fracture risk, zoledronate has been found in some studies to decrease mortality in humans and extend lifespan and healthspan in animals. Because senescent cells accumulate with aging and contribute to multiple co-morbidities, the non-skeletal actions of zoledronate could be due to senolytic (killing of senescent cells) or senomorphic (inhibition of the secretion of the senescence-associated secretory phenotype [SASP]) actions. To test this, we first performed *in vitro* senescence assays using human lung fibroblasts and DNA repair-deficient mouse embryonic fibroblasts, which demonstrated that zoledronate killed senescent cells with minimal effects on non-senescent cells. Next, in aged mice treated with zoledronate or vehicle for 8 weeks, zoledronate significantly reduced circulating SASP factors, including CCL7, IL-1β, TNFRSF1A, and TGFβ1 and improved grip strength. Analysis of publicly available RNAseq data from CD115+ (CSF1R/c-fms+) pre-osteoclastic cells isolated from mice treated with zoledronate demonstrated a significant downregulation of senescence/SASP genes (SenMayo). To establish that these cells are potential senolytic/senomorphic targets of zoledronate, we used single cell proteomic analysis (cytometry by time of flight [CyTOF]) and demonstrated that zoledronate significantly reduced the number of pre-osteoclastic (CD115+/CD3e-/Ly6G-/CD45R-) cells and decreased protein levels of p16, p21, and SASP markers in these cells without affecting other immune cell populations. Collectively, our findings demonstrate that zoledronate has senolytic effects *in vitro* and modulates senescence/SASP biomarkers *in vivo*. These data point to the need for additional studies testing zoledronate and/or other bisphosphonate derivatives for senotherapeutic efficacy.

## Introduction

Aging is a major risk factor for multiple co-morbidities, including sarcopenia [1, 2], cardiovascular disease [3], cognitive disorders [4], arthritis [5], respiratory disorders [6], osteoporosis [7], and frailty [8]. Although treatments exist to individually alleviate the symptoms and/or progression of each of these diseases, with the increasing elderly population, the overall burden of these diseases is expected to increase markedly. Indeed, the global population aged 60 years and older is expected to increase from 8.5% to 16.7% of the population by 2050, thereby outnumbering adolescents and young individuals combined (aged 10-24 years) [9]. With this expectation of such a large demographic of aged individuals, the need to study, understand, and treat age-related disorders is more important than ever.

It has been postulated that targeting the fundamental mechanisms of aging may, in fact, delay or alleviate most age-related diseases. A well-studied contributor to aging is the age-related onset of cellular senescence, which is a state of growth arrest that occurs due to the accumulation of DNA damage and cellular stress [7, 10]. This is distinct from quiescence, as senescent cells are excluded from a reversible G0 state [11] and also acquire the senescence-associated secretory phenotype (SASP), which is characterized by the release of pro-inflammatory factors that have detrimental effects both locally and systemically on tissue function [7]. In addition, these SASP factors can further induce senescence in other cells [12, 13]. Chronic accumulation of senescent cells has been linked to common age-related diseases [10], and clearance of senescent cells in mice either through pharmacological or genetic methods extends healthspan [13, 14]. Further, senescent cell clearance in mice also alleviates the progression of systemic diseases, including cardiovascular disease [15, 16], physical frailty [13], pulmonary fibrotic disease [17], cancer [18], and bone loss related to aging [19] or cancer treatments [20, 21]. However, pharmacological therapies that clear or suppress senescent cells are in very early-stage clinical trials, with no “senotherapeutics” currently available for the treatment of osteoporosis or age-related diseases in humans.

Zoledronate, a commonly prescribed bisphosphonate for patients with osteoporosis or skeletal complications due to multiple myeloma or cancer metastases, may be a potential candidate drug for targeting cellular senescence. Zoledronate has an established safety profile and has been approved for clinical use for nearly 20 years [22]. Treatment with zoledronate reduces bone turnover, improves bone mineral density (BMD), and reduces fracture risk by as much as 70% [23]. Interestingly, zoledronate treatment has also been associated in individual studies with reduced risks of mortality [24, 25], cancer [26, 27], and cardiovascular disease [27-29]; however, a recent meta-analysis of 6 clinical trials of zoledronate failed to find a significant effect of zoledronate on overall mortality, although the authors acknowledged some uncertainty regarding these findings due to significant heterogeneity of the results [30]. Specifically, 2 large clinical trials of zoledronate treatment found 28% and 35% reductions in mortality [24, 26] that were not observed in other, smaller studies. Moreover, an earlier meta-analysis had demonstrated an overall reduction in mortality with use of bisphosphonates [29], although zoledronate was not specifically analyzed in that study.

Several mechanistic animal studies have shown that zoledronate treatment has beneficial non-skeletal effects, including increasing lifespan in a mouse model of Hutchinson-Gilford progeria syndrome [31], inhibition of tumor growth and metastasis [32, 33], cellular protection against radiation [34], and improved muscle function in mice after chemotherapy [35, 36]. Interestingly, these effects in models associated with DNA damage, a robust trigger for cellular senescence, are similar to those seen after senescent cell clearance. However, it is unclear if the skeletal or extra-skeletal effects of zoledronate are related to either senolytic (*i*.*e*. killing of senescent cells) or senomorphic (*i*.*e*. inhibition of the secretion of the SASP) effects or are unrelated to the senescence pathway. Thus, in the present study, we used multiple complementary approaches (*in vitro, in vivo*, and *in silico*) to evaluate possible effects of zoledronate on modulating cellular senescence.

## Results

### Zoledronate has senolytic activity *in vitro*

To determine the effect of zoledronate on cellular senescence, human lung fibroblasts (IMR90 cells) that were induced to become senescent by etoposide were treated with increasing concentrations of zoledronate, and the percent of SA-β-gal+ senescent cells following treatment was determined by staining with the fluorogenic substrate 5-dodecanoylaminofluorescein di-β-D-galactopyranoside (C_12_FDG), as previously described [37, 38]. As shown in Fig. 1A and C, zoledronate, in a dose range that is typically used to suppress osteoclast formation *in vitro* [39], preferentially reduced the number of C_12_FDG-positive senescent cells in a dose-dependent manner. More importantly, zoledronate treatment of non-senescent, proliferating IMR90 cells exhibited minimal cytotoxicity in comparison to senescent IMR90 cells (Fig 1B), indicating that the senolytic effect of zoledronate was very specific to senescent cells. Moreover, the senolytic effect of zoledronate featured an extremely high selectivity index (SI) of 93.3, which is the ratio of half maximal effective concentration (EC_50_) values for non-senescent *vs*. senescent cells (Fig. 1B). In general, an SI value ≥ 10 identifies a compound that is worthy of further investigation [40]. These findings were extended to a second model system where DNA repair-deficient mouse embryonic fibroblasts (MEFs) from *Ercc1*-deficient (*Ercc1*^*-/-*^) mice [41] were generated. The *Ercc1*^*-/-*^ MEFs were cultured at 20% O_2_ for 3 passages to induce senescence by oxidative stress where about half of the cells became senescent [37]. As shown in Suppl. Fig. 1A-C, findings with the *Ercc1*^*-/-*^ MEFs were very similar to those with the human lung fibroblasts. Taken together, these results demonstrate that zoledronate is a potent and selective senolytic in both human and mouse cells *in vitro*.

**Figure 1.**
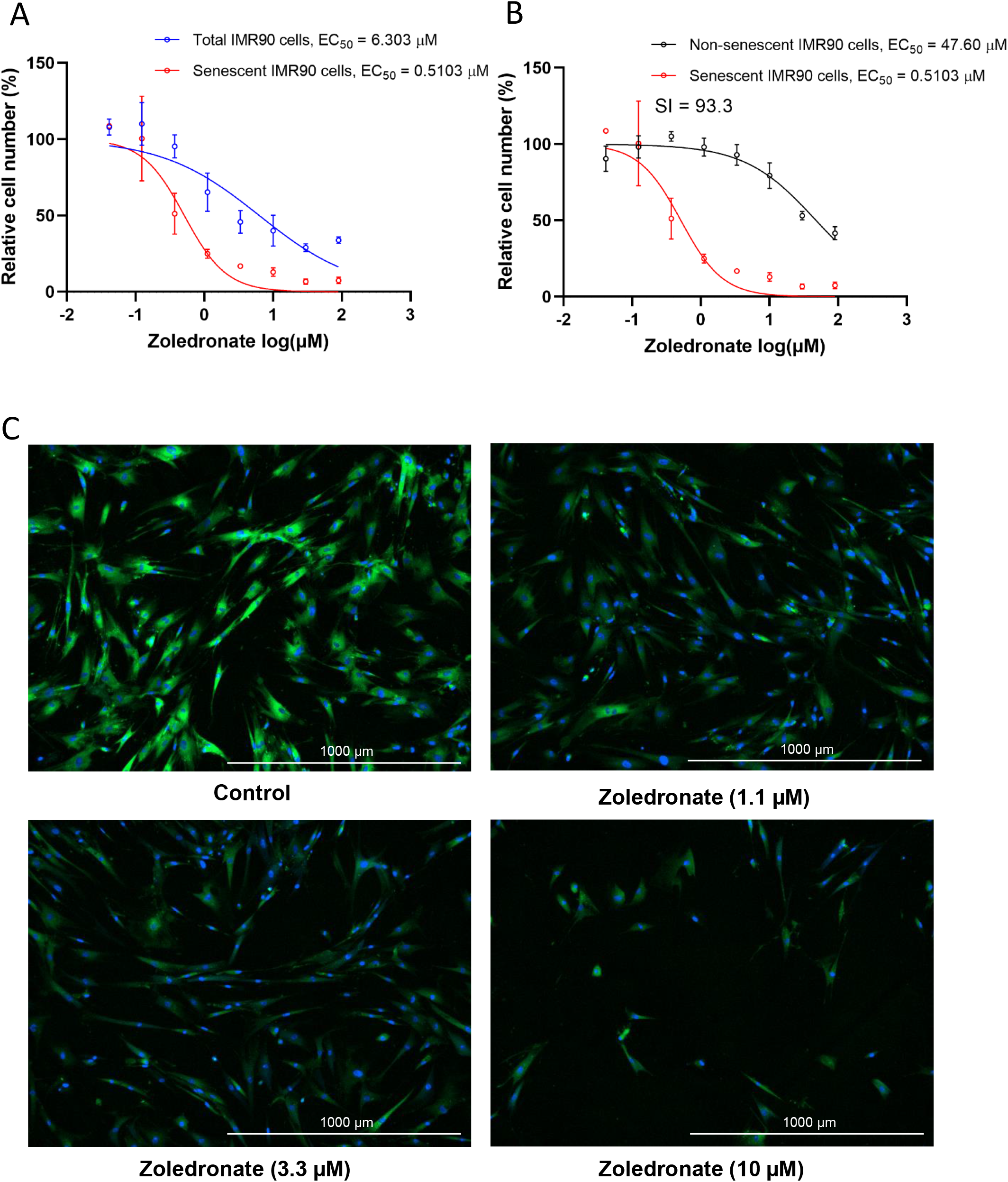
Zoledronate has senolytic effects in human lung fibroblast IMR90 cells. (A) Increasing concentrations (0.04-90 µM) of zoledronate were tested for 72 h in IMR90 cells. The figure shows the percentage of total cells (blue) and senescent IMR90 cells remaining following treatment. n = 3, mean ± SD; (B) Percentage of non-senescent (black) and senescent IMR90 cells (red) remaining after 72 hours of treatment. n = 3; (C) Representative images of C_12_FDG-based senescence assay of zoledronate in senescent IMR90 cells. Blue fluorescence indicates nuclear staining with Hoechst 33324, and bright green fluorescence indicates SA-β-gal positive senescent cells whereas dim green fluorescence represents SA-β-gal low or negative, non-senescent cells. Images were taken using Cytation 1 at 4X.

### Effects of zoledronate on circulating SASP factors and markers of frailty in aged mice

Based on the above *in vitro* data, we next evaluated whether possible senolytic effects of zoledronate would be reflected by a reduction in circulating SASP proteins *in vivo* in mice. We found that in 24 month old mice who had been treated with zoledronate for 8 weeks, there was a significant overall reduction (harmonic mean *p*-value = 0.0212, see Statistical Methods) in a panel of established SASP proteins [7, 42, 43] in zoledronate-*vs*. vehicle-treated mice (Fig. 2A), with individually significant reductions in CCL7, IL-1β, TNFRSF1A, and TGFβ1 (Fig. 2B). The potential biological consequence of the reduction in circulating SASP factors was assessed by physical function testing, which revealed that grip strength improved significantly with zoledronate as compared to vehicle treatment (Fig. 2C). However, we did not observe significant improvements in hang endurance or treadmill endurance in the zoledronate-*vs*. vehicle-treated mice (Fig. 2D, E). Findings were similar when these parameters were normalized by body weight (Fig. 2F-H). In a separate cohort treated identically (see Methods), we obtained muscle weights, which did not differ between groups (Suppl. Fig. 2A, B) as well as myofiber cross-sectional areas (CSA) at the quadriceps muscle (Suppl. Fig. 2C-E), which were also unchanged following zoledronate treatment. As shown in Suppl. Fig. 3A-C, there were no significant changes in body weight, lean mass, or fat mass in either the zoledronate- or vehicle-treated mice over the 8 weeks of treatment.

**Figure 2.**
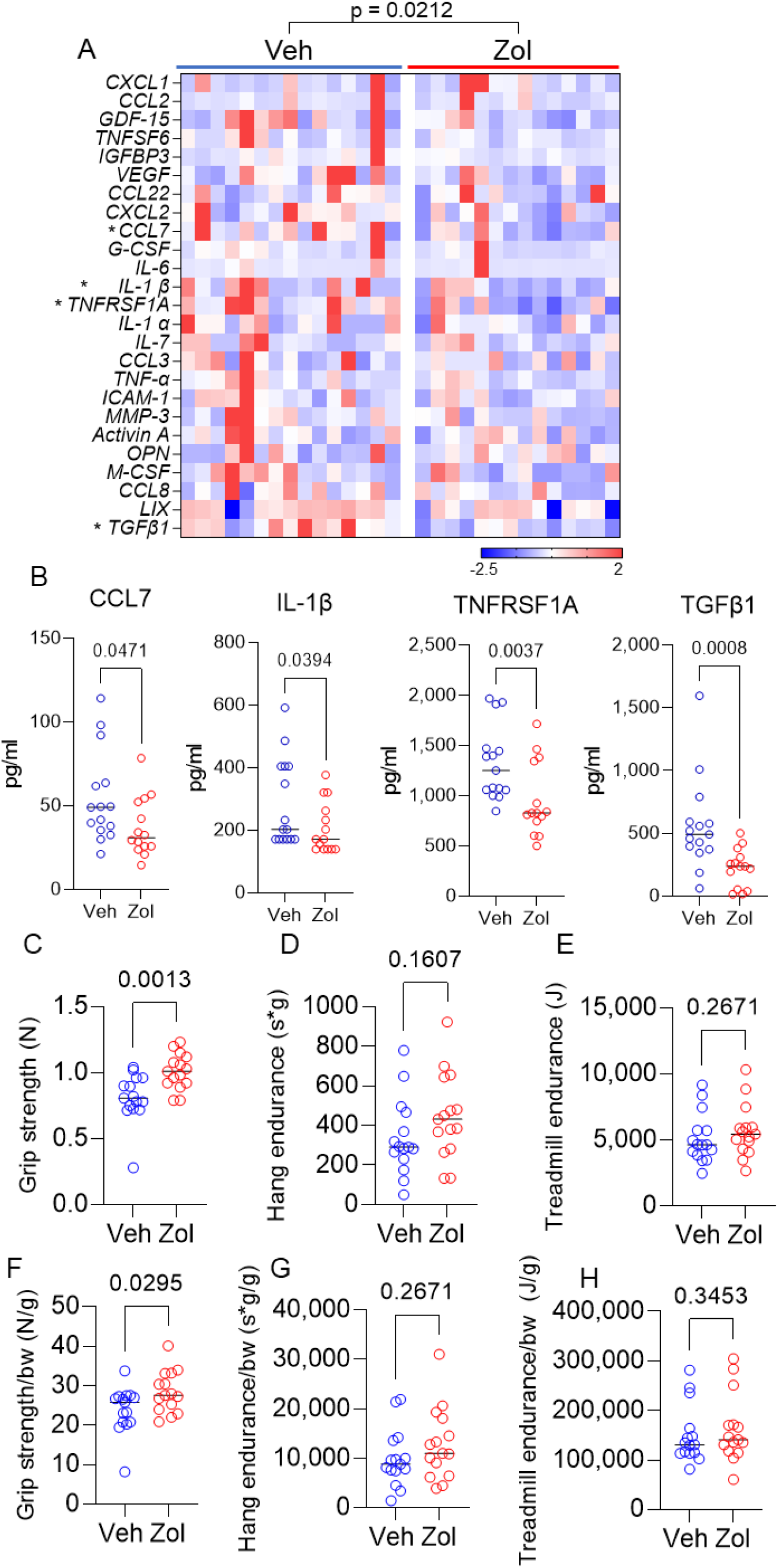
Zoledronate reduces circulating SASP markers and improves grip strength in old mice. (A) Zoledronate significantly reduces a panel of SASP markers in old mice (overall *p*-value is using the harmonic mean approach [see Methods]); (B) changes in plasma CCL7, IL1β, TNFRSF1A, and TGFβ1 in the vehicle-*vs*. zoledronate- treated mice; effects of zoledronate on (C) grip strength; (D) hang endurance; and (E) treadmill endurance. Panels F-H show these parameters normalized to body weight. *p*-values are using the Mann-Whitney test; n = 15 Veh, n=14 Zol mice/group (A-B) and n=15 per group (C-H).

**Figure 3.**
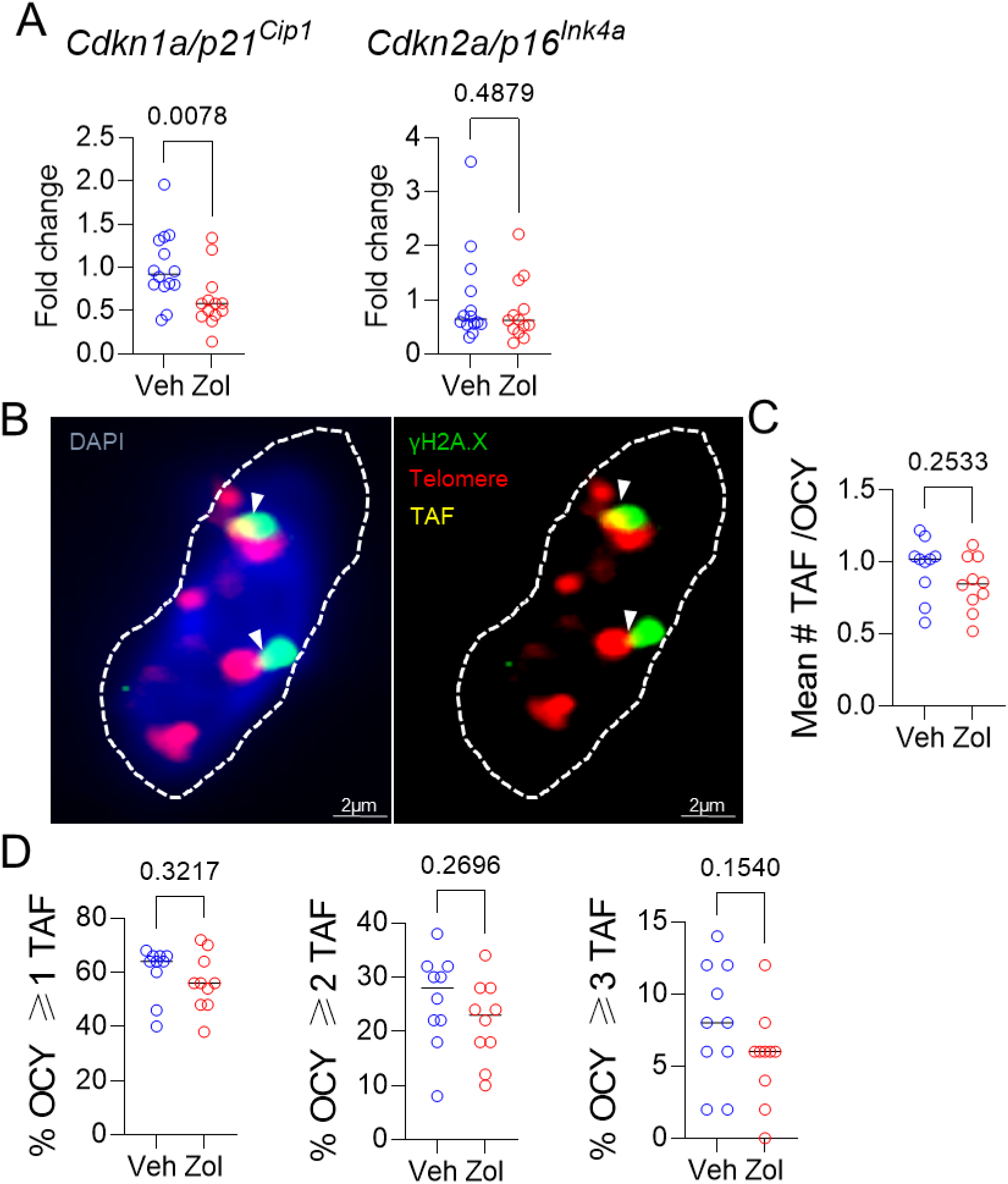
Effects of zoledronate on senescence markers in bone. (A) Zoledronate treatment led a significant downregulation of *Cdkn1a/p21*^*Cip1*^, but not *Cdkn2a/p16*^*Ink4a*^ in the centrifuged metaphyses, enriched for osteocytes (see Methods). (B) Example of an osteocyte with γH2A.X (green) and telomeres (red); where they colocalize, a TAF is scored (yellow, marked by white arrowhead). (C) Mean #TAF/osteocyte, (D) percentage of osteocytes with ≥ 1, 2, or 3 TAF/osteocyte. *p*-values are using Mann-Whitney test; n = 14 Veh and n=13 Zol mice per group (A, B) and n=10/group (B-D).

### Skeletal effects of zoledronate

As expected, spine trabecular (Suppl. Fig. 4A-D) and femur cortical (Suppl. Fig. 4E) parameters were enhanced in the zoledronate-*vs*. vehicle-treated mice. Both serum CTx (marker of bone resorption, Suppl. Fig. 4F) and P1NP (marker of bone formation, Suppl. Fig. 4G) decreased in the zoledronate-treated mice but remained unchanged in the vehicle-treated mice. As previously described in rodents treated with zoledronate [44, 45], osteoclast numbers did not differ between groups (Suppl. Fig. 4H); however, there was a marked reduction in osteoclasts that were attached to the bone surface (Suppl. Fig. 4I) and a concomitant increase in detached osteoclasts (Suppl. Fig. 4J, K), as has also been previously demonstrated in rodent models [44, 45]. Zoledronate treatment did not result in changes in osteoblast numbers (Suppl. Fig. 4L) but did reduce mineral apposition (Suppl. Fig. 4M) and bone formation (Suppl. Fig. 4N) rates, reflecting the known coupling between bone resorption and bone formation [46]. Thus, treatment with zoledronate not only reduced circulating SASP factors and improved grip strength, but also had the expected effects on improving skeletal parameters in old mice.

Given the *in vitro* senolytic effects of zoledronate as well as the significant reduction in circulating SASP factors in the zoledronate-treated mice, we next examined the bones of vehicle-*vs*. zoledronate-treated mice for differences in senescence markers. As shown in Fig. 3A, zoledronate treatment did result in a significant reduction in mRNA levels of one of the drivers of cellular senescence, *Cdkn1a/p21*^*Cip1*^, but not of *Cdkn2a/p16*^*Ink4a*^, in osteocyte-enriched bones from zoledronate-*vs*. vehicle-treated mice. Assessment of telomere-associated foci (TAF), a marker of telomeric DNA damage associated with senescence [47] (Fig. 3B), showed an overall pattern of lower mean #TAF/osteocyte (Fig. 3C) as well as the percentage of osteocytes with ≥ 1, 2, or 3 TAFs/cell (Fig. 3D), but none of these differences were statistically significant. Statistically combining these parameters in a generalized mixed effects model resulted in a treatment *p*-value for zoledronate effects on TAFs of 0.161.

### Effects of zoledronate on bone marrow pre-osteoclastic cells

As zoledronate executes its skeletal anti-resorptive actions by inhibiting bone resorption, we next investigated the potential senotherapeutic effects of zoledronate on osteoclast lineage cells. We first made use of a publicly available RNAseq dataset by Ubellacker *et al*. [48] (GSE108250) where C57BL/6J mice were treated for 5 days either with 50 μg/kg G-CSF, which is known to induce osteoclastogenesis *in vivo* [49, 50] (control), or a combination of 50 μg/kg G-CSF and 100μg/kg zoledronate. After isolation of pre-osteoclastic CD115+ (CSF1R/c-fms+) cells [51, 52] from the treated mice, RNA-sequencing of these cells was performed. Fig. 4A shows a volcano plot of the differentially regulated genes in the control *vs*. zoledronate groups. To specifically evaluate effects of zoledronate on senescence/SASP gene expression, we used a recently validated gene set from our laboratory (SenMayo) that was demonstrated to identify senescent cells across tissues and species with high fidelity [53]. As shown in Fig. 4B, this gene set was significantly downregulated in the CD115+ cells following zoledronate treatment, with similar findings noted using another publicly available SASP gene set (Fridman [54]; Fig. 4C). Additional highly downregulated gene sets by zoledronate included the inflammatory response (Fig. 4D) and IL1 signaling (Fig. 4E) pathways. These *in silico* analyses thus demonstrate that zoledronate downregulates senescence/SASP and inflammatory pathways in pre-osteoclastic cells.

**Figure 4.**
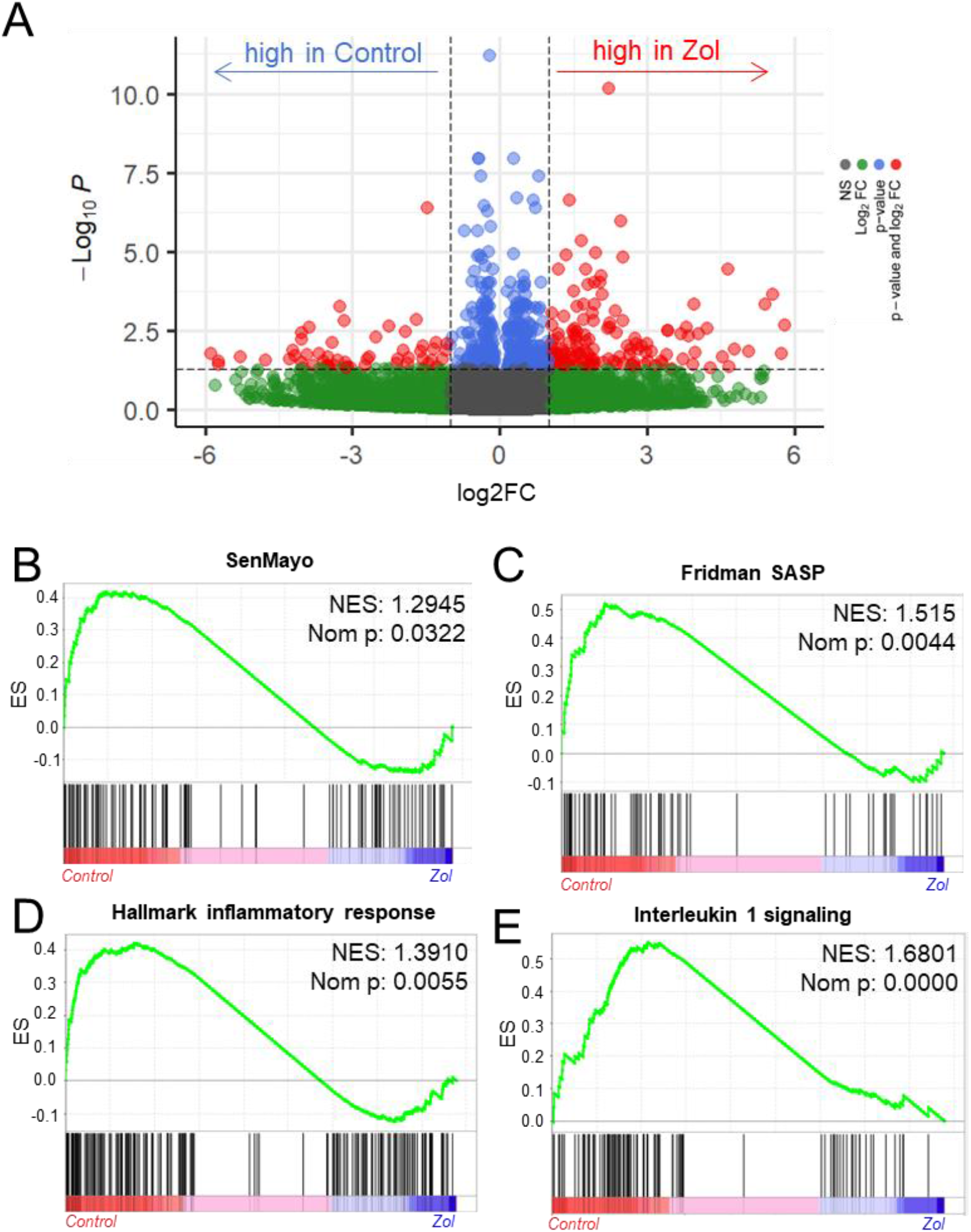
In silico analysis indicates the senolytic/senomorphic activity of zoledronate. (A) Volcano plot showing the differentially regulated genes between control and zoledronate treatments; (B) The SenMayo and (C) Fridman senescence/SASP gene sets indicate a downregulation of senescence-associated pathways in the zoledronate group, and (D) the hallmark of inflammatory response as well as (E) Interleukin 1 signaling also indicate a lower inflammatory activity in the zoledronate group. p-values are using one-way ANOVA with multiple comparisons, adjusted with the Tukey *post-hoc* method. N=8 mice in the Control and n=7 in the zoledronate group.

### Pre-osteoclastic cells are targeted by zoledronate treatment as demonstrated by cytometry by time of flight (CyTOF)

In order to extend the above RNAseq findings of Ubellacker *et al*. [48] we used a single cell proteomic approach (cytometry by time of flight, CyTOF). Suppl. Table S1 provides a list of all antibodies used in this analysis; note also that each of these antibodies have been validated for CyTOF by the Mayo Clinic CyTOF Core Laboratory, and additional validations of these antibodies in our laboratory are described in Doolittle *et al*. [55]. For these studies, we treated 21-month-old female mice with zoledronate or vehicle for two weeks. After isolation of the hematopoietic cell population, a total of 7,371,979 cells were analyzed by CyTOF. Fig. 5A shows the expression profile of the identified clusters, and the FlowSOM clustering algorithm [56] revealed one population to be high in CD115 (Fig 5A, arrow). This cluster, which was negative for T cell (CD3e), neutrophil (Ly6G), and B cell (CD45R) markers, also had the highest expression of p16, CD117 (cKit), and Cathepsin K; p21 was present but not as high as in some other cell types (*e*.*g*., neutrophils; Fig. 5B-D). Further, as shown in Fig. 5D, these CD115+ cells also expressed high levels of the DNA damage marker, phospho-ATM [57], and were the most highly SASP-positive hematopoietic cells in the aged bone marrow microenvironment, expressing high levels of IL-1α, IL-1β, CXCL1, and PAI-1. Importantly, zoledronate clearly targeted these pre-osteoclastic cells, as this population (CD115+/CD3e-/Ly6G-/CD45R-) was significantly reduced following zoledronate treatment (by 29%, Fig. 6A). This effect of zoledronate was specific for the CD115+ cells as zoledronate did not reduce the percentages of B- or T-cells, neutrophils, monocytes, or dendritic cells (Suppl. Fig. 5A-E). Moreover, the key senescence markers p16 as well as p21 were significantly downregulated in CD115+ cells, as were the SASP markers IL1-β and PAI1 (*Serpine1*) (Fig. 6B). These findings thus demonstrate that the target of zoledronate within the bone microenvironment is CD115+ cells which are reduced both in number and in their expression of key senescence/SASP markers following zoledronate treatment.

**Figure 5.**
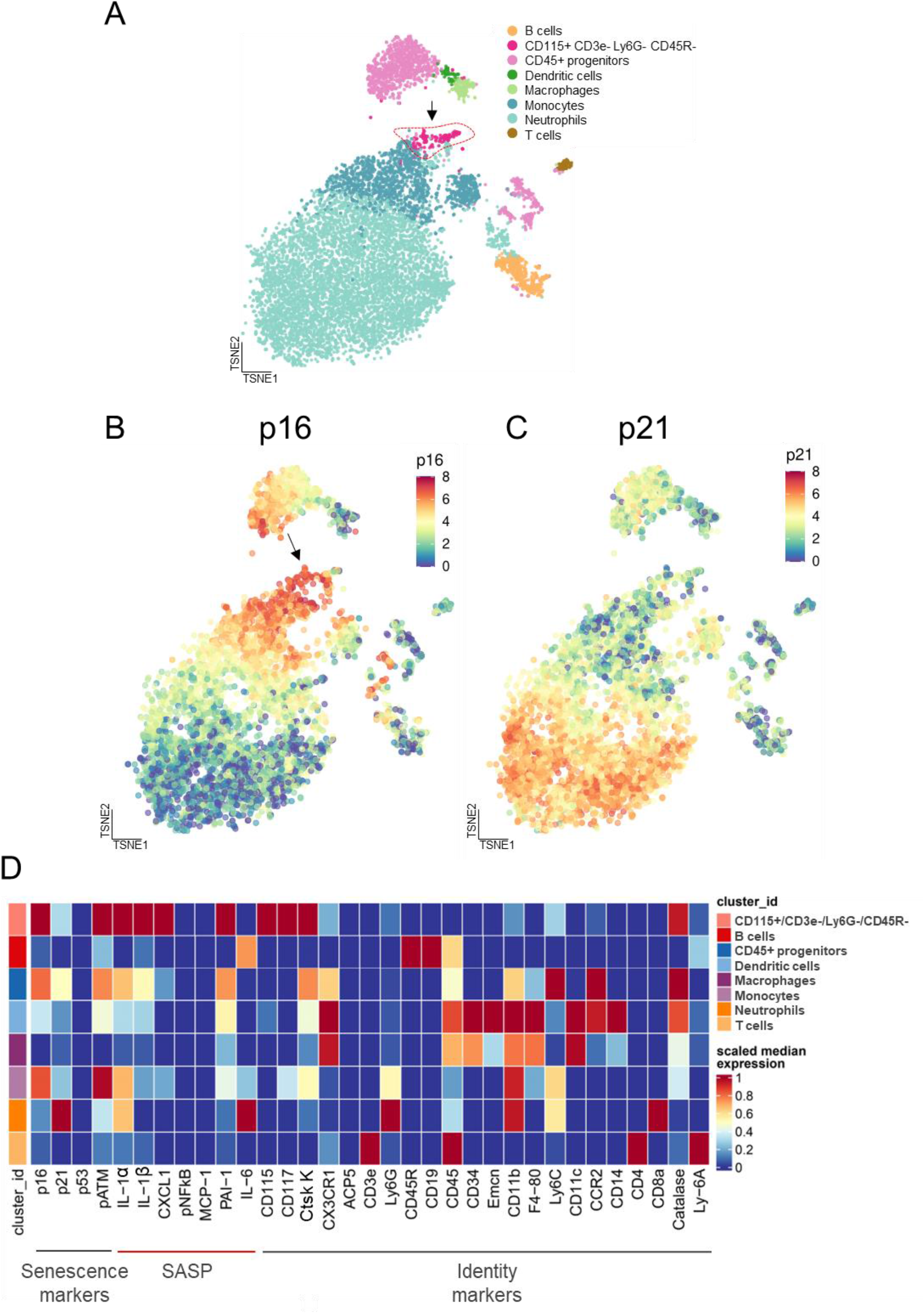
Cytometry by time of flight (CyTOF) reveals the cellular targets of zoledronate treatment. (A) CyTOF of hematopoietic cells led to distinct clusters. A t-distributed Stochastic Neighbor Embedding (tSNE) visualization of vehicle treated specimens (n=10) shows the integration of the datasets with the arrow pointing to the CD115+/CD3e-/Ly6G-/CD45R-cells; (B) p16 expression and (C) p21 expression in a tSNE visualization in the Veh group demonstrates a high expression of p16 in the CD115+ cells; (D) Heatmap showing the expression of senescence, SASP, and identity markers in the bone marrow hematopoietic cells, demonstrating that the CD115+/CD3e-/Ly6G-/CD45R-cells express the highest levels of p16, pATM (DNA damage marker), and SASP proteins.

**Figure 6.**
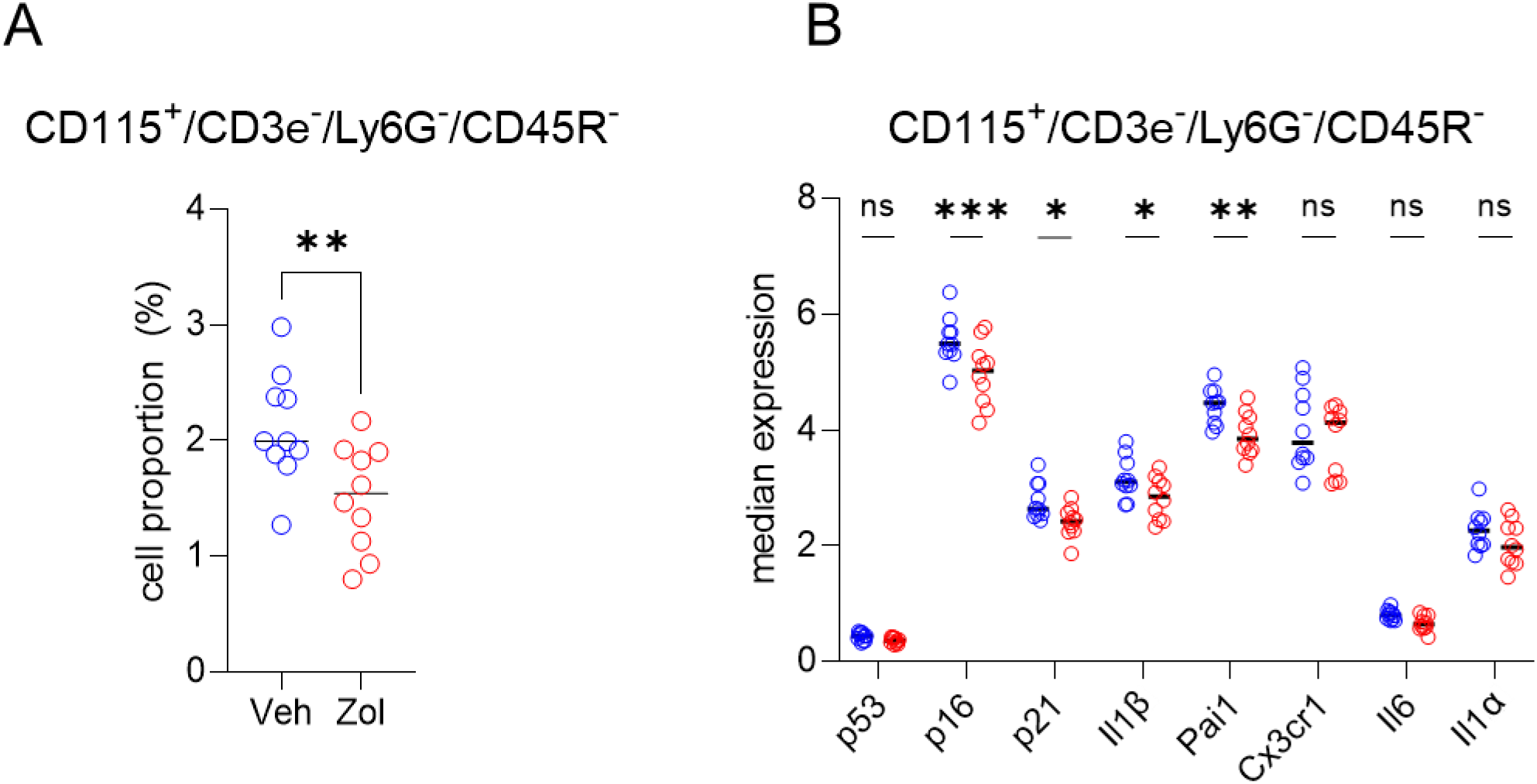
CD115^+^ cells are targeted by zoledronate. (A) Percent CD115^+^ cells is significantly reduced after zoledronate treatment (unpaired t-test, *p*<0.001): (B) The median expression in the vehicle *vs*. zoledronate samples shows a significant downregulation of p16 (*p*=0.0001), p21 (p<0.05) and the SASP markers, IL1β (p<0.05) and PAI1/SERPINE1 (*p*<0.001) (two-way ANOVA with FDR correction for multiple comparisons, method by Benjamini, Krieger and Yekutieli). N=10 mice in the vehicle and n=10 in the zoledronate group.

## Discussion

In the present study, we used multiple, complementary approaches to evaluate the possible effects of zoledronate on cellular senescence. The evidence we provide that the beneficial extra-skeletal effects of zoledronate may be mediated, at least in part, by modulation of cellular senescence is the following: (1) *In vitro*, zoledronate exhibited potent senolytic effects with a high selectivity index on both human and mouse senescent cells; (2) *in vivo*, in aged mice, treatment with zoledronate was associated with a significant reduction in a panel of circulating SASP factors concomitant with an improvement in grip strength, although we acknowledge that other markers of frailty were not improved by zoledronate; (3) we complemented these findings with *in silico* analysis of an independent study and model by Ubellacker *et al*. [48] that included RNAseq analysis of pre-osteoclastic CD115+ (CSF1R/c-fms+) cells [51, 52], which revealed a significant reduction in two panels of senescence/SASP genes (SenMayo and Fridman) in these cells by zoledronate; and (4) we directly tested for possible senolytic/senomorphic effects of zoledronate using single cell proteomic analysis (CyTOF) and demonstrated that within the hematopoietic bone microenvironment, a subset of pre-osteoclastic cells (CD115+/CD3e-/Ly6G-/CD45R-) expressed high levels of p16 and SASP markers and that these cells were specifically reduced by zoledronate, along with a decrease in the expression of several SASP markers.

Our finding that myeloid cells expressing CSF1R (CD115+) are specifically targeted by zoledronate and are the most highly enriched hematopoietic cells in senescence/SASP/inflammatory genes and proteins are of considerable interest in light of the demonstration by Ambrosi and colleagues [58] that the ligand for CSF1R, *Csf1*, was highly upregulated in skeletal stem cells from aged (24 month) compared to young (2 month) mice. Thus, the targeting of the CSF1R+ cells by zoledronate may be particularly important in terms of interrupting an inflammatory loop between aging skeletal stem cells and myeloid cells, thereby contributing to the beneficial non-skeletal effects of zoledronate.

It is of interest that, consistent with our findings showing senescent pre-osteoclasts as key targets of zoledronate, work by Su et al. [59] has identified an apparently very similar population of pre-osteoclastic cells expressing senescence/SASP markers that accumulate in subchondral bone in a mouse model of osteoarthritis in the setting of a high fat diet. In that study, inhibiting the SASP from these cells attenuated the progression of the osteoarthritis. Moreover, previous work from the same laboratory also demonstrated that aged mice fed a high fat diet had high serum PDGF-BB levels that were derived, in fact, from bone marrow pre-osteoclastic cells [60] and these investigators linked the increased PDGF-BB to bone loss as well as arterial stiffening. Thus, there is an evolving body of evidence implicating pre-osteoclastic cells expressing senescence/SASP markers as playing important roles in mediating tissue dysfunction in various settings. We should note, however, that senescence was originally described in the context of mesenchymal cell populations [61], and further studies are needed to compare features of senescence in mesenchymal cells *vs*. the apparent senescent phenotype of immune/hematopoietic cell populations.

In terms of mesenchymal cells, we did demonstrate potent senolytic effects of zoledronate on senescent human lung and mouse embryonic fibroblast cells *in vitro. In vivo*, zoledronate treatment reduced *Cdkn1a/p21*^*Cip1*^, but not *Cdkn2a/p16*^*Ink4a*^, mRNA levels in osteocyte-enriched bone fractions from zoledronate-compared to vehicle-treated mice. Using the highly specific TAF assay for telomeric DNA damage, we did observe a trend for reduced TAFs in osteocytes with zoledronate treatment, but the differences did not reach statistical significance. Given the labor-intensive nature and variability of this assay, it is possible that a larger sample size would have resulted in a significant reduction in osteocytic TAFs in the zoledronate-treated mice. Thus, although our findings clearly implicate myeloid CD115+ cells as *bona fide* targets of the senotherapeutic and/or anti-inflammatory effects of zoledronate *in vivo*, it is possible that zoledronate also targets senescent mesenchymal cells, and further studies are needed define the senescent cell populations *in vivo* more clearly that may be reduced by zoledronate.

Our findings in mice are also consistent with recent findings in *Drosophila* demonstrating that zoledronate treatment was associated with an extension of lifespan and improved climbing ability, which is analogous to assessment of grip strength in mice [62]. This study also linked the known inhibition of farnesyl pyrophosphate synthetase (FPPS) by zoledronate to a reduction in the accumulation of DNA damage, which is a key trigger of cellular senescence [10]. These findings in *Drosophila* are similar to previous work from the same group demonstrating that zoledronate extended the lifespan of human bone marrow stromal cells (BMSCs) *in vitro*, and this was associated with a reduction in markers of DNA damage as well as reduced p21 and p16 protein expression [34].

Although our studies, when combined with the previous work noted above, point to potential effects of zoledronate on modulating cellular senescence and the SASP, we recognize important limitations. Specifically, although in this “proof-of-concept” study we administered zoledronate twice weekly to mice using a well-established regimen for rodent studies [63], patients with osteoporosis are generally only treated with zoledronate once annually. However, it is possible that ongoing release of zoledronate from the skeleton, as has been demonstrated to occur [64], may not only inhibit osteoclastic bone resorption, but also affect the highly inflammatory CD115+ cells we describe here that are also present in the bone microenvironment and reduce both their number and secretion of SASP/inflammatory factors, thereby leading to beneficial physiological effects. Clearly, further studies are needed to address this possibility. We also note that due to cost and availability of aged mice, we only studied female mice; while this is consistent with the epidemiology of osteoporosis in that fractures in women far outnumber those in men [65] and thus far more women than men are treated with zoledronate, additional studies are needed to evaluate possible sex differences in the effects of zoledronate on senescence and SASP markers we observed. These limitations notwithstanding, given our *in vitro* data of senolytic effects of zoledronate and corroboratory *in vivo* findings, our work should provide an impetus to develop zoledronate (or other bisphosphonate) analogs with greater senolytic efficacy as well as distribution to non-skeletal tissues for beneficial effects not only in bone, but across multiple aging organs.

## Methods

### Cell culture

Human lung fibroblast IMR90 cells were obtained from American Type Culture Collection (ATCC) and cultured in EMEM medium supplemented with 10% fetal bovine serum. To induce senescence, IMR90 cells were treated with 20 μM etoposide for 24 h. After etoposide removal, cells were cultured in normal medium for additional 5 days before being collected for senescence assay. Primary *Ercc1*^*-/-*^ mouse embryonic fibroblasts (MEFs) were isolated on embryonic day 12.5-13.5. In brief, mouse embryos were isolated from yolk sac followed by the complete removal of viscera, lung, and heart if present. Embryos were then minced into fine chunks, fed with MEFs medium, and cultivated under 3% O_2_ to reduce stresses. Cells were split at 1:3 when reaching confluence. MEFs were grown at a 1:1 ratio of Dulbecco’s Modification of Eagles Medium (supplemented with 4.5 g/L glucose and L-glutamine) and Ham’s F10 medium, supplemented with 10% fetal bovine serum, penicillin, streptomycin, and non-essential amino acid. To induce oxidative stress-mediated DNA damage, *Ercc1*^*-/-*^ MEFs were switched to 20% O_2_ for 3 passages [37].

### SA-β-gal senescence assay by C_12_FDG staining

Senescent IMR90 cells or *Ercc1*^*−/−*^ MEFs were seeded at 2000 cells per well in black walled, clear bottomed 96-well plates at least 6 h prior to treatment. Following the addition of zoledronate or control, the IMR90 cells were incubated for 72 h and the *Ercc1*^*−/−*^ MEFs for 48 h at 20% O_2_. After removing the medium, cells were incubated in 100 nM Bafilomycin A1 in culture medium for 1 h to induce lysosomal alkalinization. Then 10 μM or 20 μM of fluorogenic substrate C_12_FDG (Setareh Biotech, USA) were added to IMR90 cells or MEFs, respectively, and incubated for 2 h, followed by counterstaining with 2 μg/ml Hoechst 33342 (Thermo Fisher, USA) for 15 min. Subsequently cells were washed with PBS and fixed in 2% paraformaldehyde for 15 min. Finally, cells were imaged with 6 fields/well using a high content fluorescent image acquisition and analysis platform Cytation 1 (BioTek, VT, USA) [37, 38].

### Study Approval

Animal studies were performed under protocols approved by the Institutional Animal Care and Use Committee (IACUC) and experiments were performed in accordance with Mayo Clinic IACUC guidelines.

### Animal studies

The study included three mouse cohorts. Note that in order to maximize power and given that zoledronate is used in a far greater proportion of women than men, we only studied female mice. The first cohort included 22-month-old female *C57BL/6N* mice (n=15/group) that were treated intraperitoneally with zoledronate (125 µg/kg) or vehicle twice weekly for 8 weeks, using a regimen established by Dr. Russell and the Sheffield group [63]. In this cohort we assessed body composition, circulatory SASP factors, bone turnover markers, frailty, skeletal µCT analysis, qPCR analysis of metaphyseal bone samples, and TAFs. The second cohort was to assess muscle weights and muscle fiber cross-sectional area. In this cohort, 21-month-old female *C57BL/6N* mice (n=10/group) were treated intraperitoneally with the same zoledronate regimen (125 µg/kg twice weekly) or vehicle for 8 weeks. Finally, for the cytometry by time of flight (CyTOF) studies, mice were treated for two weeks with either zoledronate (n=10) or vehicle (n=10), the mice were sacrificed, and one mouse tibia isolated. All mice were housed in ventilated cages in a pathogen-free, accredited facility under a 12-hour light/12-hour dark cycle with constant temperature (23°C) and access to food and water *ad libitum*. All assessments were performed in a blinded fashion.

### Assessment of body composition

Body mass (grams) was recorded on all mice at the onset of the study and 2, 4, and 6 weeks after zoledronate or vehicle treatment and finally at the endpoint (week 8). Body composition (*i*.*e*. whole-body lean grams and fat mass grams) was assessed at the baseline and endpoint of the study *in vivo* in non-anesthetized and conscious mice using quantitative Echo magnetic resonance imaging (EchoMRI-100), as described [66].

### Mouse tissue collections

Prior to sacrifice, body mass was recorded in the morning (under nonfasting conditions) and serum/plasma was collected from anesthetized mice by cardiac puncture at time of death and stored at –80 °C. After euthanasia, the tibiae, femurs, humeri, and vertebrae were excised from the mice and skeletal muscle/connective tissues were removed. The right femur and part of the lumbar vertebrae (L4–6) were fixed in ethanol (EtOH) for *ex vivo* μCT scanning followed by histomorphometry. Another part of the lumbar vertebrae (L1-3) were embedded in methyl methacrylate and sectioned for TAFs. The metaphysis of the R tibia was cut and centrifuged to remove bone marrow and obtain an osteocyte-enriched sample for further analysis, as previously described [67]. The samples were immediately homogenized in QIAzol Lysis Reagent (QIAGEN) and frozen at –80°C for rt-qPCR mRNA gene expression analyses.

### Measurement of circulating SASP factors

The concentrations of CCL2, CCL3, CCL7, CCL8, CCL22, CXCL1, CXCL2, G-CSF, GDF-15, ICAM-1, IGFBP3, IL-1α, IL-1β, IL-7, IL-17, LIX, M-CSF, MMP-3, OPN, TNFSF6, TNFRSF1A, TNF-α, and VEGF in plasma were quantified using commercially available multiplex magnetic bead immunoassays (R&D Systems) based on the Luminex xMAP multianalyte profiling platform and analyzed on a MAGPIX System (Merck Millipore). All assays were performed according to the manufacturer’s protocols. Activin A and TGF-β1 concentration were determined by a Quantikine ELISA Kit (R&D Systems) according to the manufacturer’s instructions. Investigators were blinded as to sample identity and treatment of mice.

### Assessment of physical function

All measurements were performed at least 5 days after the last dose of zoledronate or vehicle treatment as previously described [13]. Forelimb grip strength (N) was determined using a Grip Strength Meter (Columbus Instruments, Columbus, OH). Results were averaged over ten trials. For the hanging test, mice were placed onto a 2-mm-thick metal wire that was 35 cm above a padded surface. Mice were allowed to grab the wire with their forelimbs only. Hanging time was normalized to body weight as hanging duration (sec) × body weight (g). Results were averaged from 3 trials for each mouse. For treadmill performance, mice were acclimated to a motorized treadmill at an incline of 5° (Columbus Instruments) over 3 days for 5 min each day, starting at a speed of 5 m/min for 2 min and progressing to 7 m/min for 2 min and then 9 m/min for 1 min. On the test day, mice ran on the treadmill at an initial speed of 5 m/min for 2 min, and then the speed was increased by 2 m/min every 2 min until the mice were exhausted. Exhaustion was defined as the inability to return onto the treadmill despite a mild electrical shock stimulus and mechanical prodding. Distance was recorded and total work (kJ) was calculated using the following formula: mass (kg) × g (9.8 m/s^2^) × distance (m) × sin(5°).

### Evaluation of myofiber cross-sectional areas

Quadriceps muscles were embedded in Optimal Cutting Temperature compound (Sakura Finetek, Torrance, CA, USA), frozen in liquid nitrogen-cooled 2-methylbutane (Sigma, St. Louis, MO, USA), and stored at -80°C. Transverse 7 μm-thick frozen sections were cut with a Leica cryostat and mounted onto SuperFrost Plus slides (Thermo Fisher Scientific, Waltham, MA, USA). Muscle sections were dried for 2 hours at -20°C, and then stored at -80°C until analysis. Frozen quadriceps sections were removed from -80°C and fixed in 4% paraformaldehyde (PFA) for 15 minutes. After being permeabilized and blocked, the sections were incubated with rabbit anti-laminin antibody (L9393, Sigma-Aldrich) overnight at 4°C, followed by goat-anti-rabbit cross-adsorbed secondary antibody Alexa Fluor 488 (A11008, Invitrogen by Thermo Fisher Scientific) for 1 hour in a humidified chamber at room temperature. Sections were mounted using ProLong Gold Antifade Mountant with DAPI (Invitrogen by Thermo Fisher Scientific). Images were captured at 10x magnification with a Zeiss Axio Imager microscope. Myofiber cross-sectional area, demarcated by Laminin staining, was quantified using MuscleJ[68].

### Skeletal µCT analysis

All imaging and analysis were performed in a blinded fashion as described by our group previously [19, 42]. At study endpoint, *ex vivo* quantitative analyses of bone microarchitecture of the lumbar vertebrae (L5) and right femur (mid-shaft diaphysis) were performed. Scan settings were as follows: 55 kVp, 10.5 μm voxel size, 21.5 diameter, 145mA, 300 ms integration time. μCT parameters were derived using the manufacturer’s protocols. Trabecular bone parameters were at the lumbar spine (200 slices). Furthermore, at the mid-diaphysis (50 slices) of the right femur, cortical thickness (Ct.Th; mm) was assessed.

### Bone turnover marker measurements

All biochemical assays for bone turnover markers (P1NP and CTX) were performed in blinded fashion. At the study endpoint, serum was collected in the morning (under non-fasting conditions) from anesthetized mice by cardiac puncture and stored at -80°C in aliquots for additional biochemical assays, which included circulating levels of bone turnover markers. Specifically, the serum bone formation marker P1NP (amino-terminal propeptide of type I collagen; ng/mL) was measured using the rat/mouse P1NP enzyme immunoassay (EIA) kit (interassay coefficient of variation [CV] <10%), and the serum bone resorption marker CTx (cross-linked C-telopeptide of type I collagen; ng/mL) was measured using a RatLaps Rat/Mouse CTx EIA kit (interassay CV <10%). Kits were purchased from Immuno Diagnostic Systems (IDS, Scottsdale, AZ).

### Bone histomorphometry

All histomorphometric analyses were performed in a blinded manner. For dynamic histomorphometry, mice were injected subcutaneously with Alizarin Red (0.1mL/animal, 7.5mg/mL) and calcein (0.1 mL/animal, 2.5mg/mL) on days 9 and 2, respectively, before euthanasia. The lumbar vertebrae were isolated from WT mice treated with vehicle or zoledronate. The vertebrae were embedded in MMA, sectioned, and stained with Masson Trichrome to assess osteoblast numbers/bone perimeter (N.Ob/B.Pm,/mm), or stained for tartrate-resistant acid phosphatase (TRAP) activity to assess osteoclast numbers per bone perimeter, N.Oc/B.Pm,/mm). Alternatively, sections were left unstained to quantify trabecular mineralizing surfaces (mineral apposition rate, MAR, µ/d; bone formation rate/bone surface, BFR/BS, µm^3^/µm^2^/d), which were 100µm apart from the growth plate and 100 µm apart from the cortical anterior or posterior vertebral body perimeter. Osteoblast (N.Ob/B.Pm) and osteoclast (N.Oc/B.Pm) numbers were assessed at the same distance from cortical bone to verify trabecular assessments. All histomorphometric measurements and calculations were performed with the Osteomeasure Analysis system (Osteometrics, Atlanta, Georgia).

### rt-qPCR analysis of metaphyseal bone

Osteocyte-enriched cell preparations were generated as described in *Mouse tissue collections*, immediately homogenized in QIAzol Lysis Reagent (QIAGEN, Valencia, CA), and stored at -80°C for subsequent RNA extraction, cDNA synthesis, and targeted gene expression measurements of mRNA levels by rt-qPCR, as described [69]. Total RNA was extracted according to the manufacturer’s instructions using QIAzol Lysis Reagent followed by purification with RNeasy Mini Columns (QIAGEN, Valencia, CA). On-column RNase-free DNase solution (QIAGEN, Valencia, CA) was then applied to degrade potentially contaminating genomic DNA. RNA purity/quantity was confirmed by Nanodrop spectrophotometry (Thermo Fisher Scientific, Wilmington, DE). Standard reverse transcriptase was performed using the High-Capacity cDNA Reverse Transcription Kit (Applied Biosystems by Life Technologies, Foster City, CA). Transcript mRNA levels were determined by rt-qPCR on the ABI Prism 7900HT Real Time System (Applied Biosystems, Carlsbad, CA) using murine TaqMan assays (Thermo Fisher Scientific, Wilmington, DE) to measure *p16*^*Ink4a*^ (*Cdkn2a*) and *p21*^*Cip1*^ (*Cdkn1a*), as described[69].

### Assessment of TAFs

To measure cellular senescence in vertebral trabecular bone, the TAF assay (*n* = 10/group) was performed on murine vertebral trabecular bone nondecalcified methyl methacrylate–embedded sections. Our protocol was adapted from a previous study[47]. Bone sections were deplasticized and hydrated in EtOH gradient followed by water and PBS. Antigen was retrieved by incubation in Tris-EDTA (pH 9.0) at 95°C for 15 minutes. After cooldown and hydration with water and PBS (0.5% Tween-20/0.1% Triton X-100), slides were placed in blocking buffer (1:60 normal goat serum; Vector Laboratories; S-1000; in 0.1% BSA/PBS) for 30 minutes at room temperature (RT). Primary antibody γ-H2AX (1:200; anti–γ-H2A.X rabbit monoclonal antibody, Cell Signaling Technology; 9718) was diluted in blocking buffer and incubated overnight at 4°C. The next day, slides were washed with PBS (0.5% Tween-20/0.1% Triton X-100) followed by PBS alone and then incubated for 30 minutes with secondary goat, anti-rabbit antibody biotinylated (1:200; Vector Laboratories; BA-1000) in blocking buffer. Subsequently, slides were washed with PBS (0.5% Tween-20/0.1% Triton X-100) followed by PBS alone, then incubated for 60 minutes with tertiary antibody (1:500; Cy5 Streptavidin, Vector Laboratories; SA-1500) in PBS. Slides were then washed 3 times with PBS, followed by FISH for TAF detection. Briefly, following 4% paraformaldehyde cross-linking for 20 minutes, sections were washed 3 times (5 minutes each in PBS) and dehydrated in graded (70%, 90%, and 100%, 3 minutes each) ice-cold EtOH.

Sections were then dried and denatured for 10 minutes at 80°C in hybridization buffer: 0.1 M Tris (pH 7.2), 25 mM MgCl2, 70% formamide (MilliporeSigma), 5% blocking reagent (Roche), with 1.0 μg/mL of Cy3-labeled telomere-specific (CCCTAA) peptide nucleic acid probe (TelC-Cy3, Panagene *Inc*.; F1002), followed by humidified dark room hybridization for 2 hours at RT. Sections were then washed and mounted with VECTASHIELD DAPI-containing mounting medium (Life Technologies) before image acquisition and analysis. The number of TAF/osteocyte was quantified in a blinded fashion by examining overlap of signals from the telomere probe with γ-H2AX (*i*.*e*. phosphorylation of the C-terminal end of histone H2A.X — a marker of double-strand breaks in DNA). The mean number of TAF/osteocyte in vertebral trabecular bone was quantified using FIJI (an ImageJ distribution software; NIH, https://imagej.nih.gov/ij/), and the percentage of TAF^+^ OCYs was calculated for each mouse based on the following criteria: percentage of OCYs with ≥1 TAF, percentage of OCYs with ≥2 TAF, and percentage of OCYs with ≥3 TAF.

### *In silico* analysis of zoledronate effects on pre-osteoclastic cells

For this analysis, we used publically available data from Ubellacker *et al*. [48] (GSE108250). Briefly, in those studies, transcriptome-wide gene expression data were generated from CD115^+^ (c-fms+) bone marrow cells (from femora and tibiae) from six to seven-week-old female C57BL/6J mice, treated with a single dose of 100μg/kg (i.p.) zoledronic acid (Novartis Pharmaceuticals) and/or 50 μg/kg (i.p.) recombinant human granulocyte-colony stimulating factor (G-CSF; carrier-free, BioLegend #578604) for three days. Sequencing was performed on a HiSeq2500 (Illumina^®^), fastq files were mapped to the murine reference genome mm10, and analysis was performed as previously described [70]. Significantly differentially regulated genes were selected by a Benjamini–Hochberg adjusted p-value ≤ 0.05 and log_2_-fold changes above 0.1 or below -0.1. Gene Set Enrichment Analysis (GSEA) [71, 72] was performed with default settings (1000 permutations for gene sets, Signal2Noise metric for ranking genes). Comparisons were made between OC precursors in combination therapy (zoledronate +G-CSF, n=7 and G-CSF alone, n=8). The analyses we present are new, and we focused on pathways and differentially expressed genes different from the ones in Uebellacker *et al*. [48] (GSE108250).

### Cytometry by time of flight (CyTOF)

After a two-week treatment with either zoledronate (n=10) or vehicle (n=10), the mice were sacrificed and one mouse tibia isolated. The epiphyseal region was removed, and the bone centrifuged. The resuspended bone marrow was RBC lysed, resuspended, and kept on ice. For labeling, the cells were incubated at 4°C with metal-conjugated antibodies (see. Suppl. Table 1). The cells were incubated with cell surface targeting antibodies for 45 min. Subsequently, they were fixed with 2% paraformaldehyde, washed, and incubated with antibodies for intracellular antigens for 45 min. After labeling, the cells were assayed on a Helios II mass cytometer (Fluidigm, South San Francisco, CA).

The generated Fcs files were normalized and debarcoded with Cytobank (Beckman Coulter Life Sciences, IN, US). A manual selection of alive singlet cells was performed, and the resulting fcs files uploaded in R version 4.0.2 (The R Project for Statistical Computing). The subsequent analyses have been done with the packages HDCytoData (1.10.0), flowWorkspace (4.2.0), openCyto (2.2.0), CATALYST (1.14.1) and SingleCellExperiment (1.12.0), following the vignette provided by Nowicka *et al*. [73]. The clustering followed the FlowSOM recommendations with a maximum of 20 clusters according to the “type” markers (see Suppl. Table 1).

Metal-conjugated antibodies used in this study are summarized in the Suppl. Table 1. Except for commercially available pre-conjugated antibodies (Fluidigm Sciences), all antibodies were conjugated to isotopically enriched lanthanide metals using the MaxPAR antibody conjugation kit (Fluidigm Sciences), according to the manufacturer’s recommended protocol. Labeled antibodies were stored at 4°C in PBS supplemented with glycerol, 0.05% BSA, and 0.05% sodium azide. All antibodies were tested with control beads as well as positive and negative control cells. Note that each of these antibodies have been validated for CyTOF by the Mayo Clinic CyTOF Core Laboratory; additional validations for the antibodies used are provided in Doolittle *et al*. [55].

### Statistical analyses

Graphical data are shown as medians with raw data points unless otherwise specified. The sample sizes were determined based on previously conducted and published experiments [19] in which statistically significant differences were observed among various bone parameters in response to multiple interventions in our laboratory. The animal numbers used are indicated in the figure legends; all samples presented represent biological replicates. The only samples excluded from analysis were the SASP factors from one mouse in the zoledronate group who uniformly had extremely high levels across multiple SASP factors, with many > 3 SDs beyond the mean of the other mice, indicating a systemic biological (*i*.*e*., unrecognized illness) or technical issue with the samples from that mouse. In agreement with our statistician (E.A.), this mouse was thus excluded from the SASP assays. The distribution of the data were examined using dot plots and histograms. Group comparisons were made using the non-parametric Mann-Whitney U test. The harmonic mean test [74] was used to combine the p-values from individual markers. The statistical analyses were performed using either GraphPad Prism (GraphPad Software, *Inc*., Version 9.0), R version 4.0.2 (The R Project for Statistical Computing), SPSS (IBM, Version 25) and GSEA (Broad Institute, V 4.1.0). Used R packages were EnhancedVolcano (1.10.0), gprofiler2 (0.2.0), clusterProfiler (version 3.10.1), and enrichplot (1.11.3). A *p*-value <0.05 (two-tailed) was considered statistically significant.

## Author contributions

P.D.R., P.S., and S.K. conceived and directed the project. P.S., S.K., and D.S., as well as P.D.R. and L.Z. designed the experiments and interpreted the data. Experiments were performed by S.K., P.S. D.S., L.Z., J.S., and J.K. S.K., P.S., M.L.D., and D.S. wrote the manuscript. All authors contributed ideas and reviewed the manuscript. S.K. oversaw all experimental design, data analyses, and manuscript preparation. P.S., D.S., P.D.R., and S.K. accept responsibility for the integrity of the data analysis.

## Acknowledgements

This work was supported by the German Research Foundation (DFG, 413501650) (D.S.), National Institutes of Health (NIH) grants P01 AG062413 (S.K., J.N.F., J.L.K), R01 AG076515 (S.K., D.G.M.), R21 AG065868 (S.K., J.N.F), R01 DK128552 (J.N.F.), R01 AG055529 (N.K.L.), Glenn Foundation for Medical Research (Z.A., N.K.L.), R37 AG13925 (J.L.K.), and the Connor Fund (J.L.K., T.T.), Robert J. and Theresa W. Ryan (J.L.K., T.T.), the Noaber Foundation (J.L.K., T.T.), and UKRI-Research England’s Connecting Capability Fund through the UK SPINE programme (J.R.E).

## Data Availability

All source data will be provided in the event of acceptance in principle.

## Code Availability

All relevant code information will be provided in the event of acceptance in principle.

## References

1. Cruz-Jentoft AJ, Landi F, Schneider SM, Zúñiga C, Arai H, Boirie Y, Chen LK, Fielding RA, Martin FC, Michel JP, Sieber C, Stout JR, Studenski SA, et al. Prevalence of and interventions for sarcopenia in ageing adults: a systematic review. Report of the International Sarcopenia Initiative (EWGSOP and IWGS). Age Ageing. 2014; 43(6):748–759.

2. Walston JD. Sarcopenia in older adults. Curr Opin Rheumatol. 2012; 24(6):623–627.

3. North BJ and Sinclair DA. The intersection between aging and cardiovascular disease. Circ Res. 2012; 110(8):1097–1108.

4. Murman DL. The Impact of Age on Cognition. Semin Hear. 2015; 36(3):111–121.

5. Lawrence RC, Felson DT, Helmick CG, Arnold LM, Choi H, Deyo RA, Gabriel S, Hirsch R, Hochberg MC, Hunder GG, Jordan JM, Katz JN, Kremers HM, et al. Estimates of the prevalence of arthritis and other rheumatic conditions in the United States. Part II. Arthritis Rheum. 2008; 58(1):26–35.

6. Lowery EM, Brubaker AL, Kuhlmann E and Kovacs EJ. The aging lung. Clin Interv Aging. 2013; 8:1489–1496.

7. Khosla S, Farr JN, Tchkonia T and Kirkland JL. The role of cellular senescence in ageing and endocrine disease. Nat Rev Endocrinol. 2020; 16(5):263–275.

8. Collard RM, Boter H, Schoevers RA and Oude Voshaar RC. Prevalence of frailty in community-dwelling older persons: a systematic review. J Am Geriatr Soc. 2012; 60(8):1487–1492.

9. Briggs AM, Cross MJ, Hoy DG, Sànchez-Riera L, Blyth FM, Woolf AD and March L. Musculoskeletal Health Conditions Represent a Global Threat to Healthy Aging: A Report for the 2015 World Health Organization World Report on Ageing and Health. Gerontologist. 2016; 56 Suppl 2:S243–255.

10. Tchkonia T, Zhu Y, van Deursen J, Campisi J and Kirkland JL. Cellular senescence and the senescent secretory phenotype: therapeutic opportunities. J Clin Invest. 2013; 123(3):966–972.

11. Pignolo RJ, Martin BG, Horton JH, Kalbach AN and Cristofalo VJ. The pathway of cell senescence: WI-38 cells arrest in late G1 and are unable to traverse the cell cycle from a true G0 state. Exp Gerontol. 1998; 33(1-2):67–80.

12. Nelson G, Wordsworth J, Wang C, Jurk D, Lawless C, Martin-Ruiz C and von Zglinicki T. A senescent cell bystander effect: senescence-induced senescence. Aging Cell. 2012; 11(2):345–349.

13. Xu M, Pirtskhalava T, Farr JN, Weigand BM, Palmer AK, Weivoda MM, Inman CL, Ogrodnik MB, Hachfeld CM, Fraser DG, Onken JL, Johnson KO, Verzosa GC, et al. Senolytics improve physical function and increase lifespan in old age. Nat Med. 2018; 24(8):1246–1256.

14. Baker DJ, Wijshake T, Tchkonia T, LeBrasseur NK, Childs BG, van de Sluis B, Kirkland JL and van Deursen JM. Clearance of p16Ink4a-positive senescent cells delays ageing-associated disorders. Nature. 2011; 479(7372):232–236.

15. Childs BG, Baker DJ, Wijshake T, Conover CA, Campisi J and van Deursen JM. Senescent intimal foam cells are deleterious at all stages of atherosclerosis. Science. 2016; 354(6311):472–477.

16. Roos CM, Zhang B, Palmer AK, Ogrodnik MB, Pirtskhalava T, Thalji NM, Hagler M, Jurk D, Smith LA, Casaclang-Verzosa G, Zhu Y, Schafer MJ, Tchkonia T, et al. Chronic senolytic treatment alleviates established vasomotor dysfunction in aged or atherosclerotic mice. Aging Cell. 2016; 15(5):973–977.

17. Schafer MJ, White TA, Iijima K, Haak AJ, Ligresti G, Atkinson EJ, Oberg AL, Birch J, Salmonowicz H, Zhu Y, Mazula DL, Brooks RW, Fuhrmann-Stroissnigg H, et al. Cellular senescence mediates fibrotic pulmonary disease. Nat Commun. 2017; 8:14532.

18. Alimirah F, Pulido T, Valdovinos A, Alptekin S, Chang E, Jones E, Diaz DA, Flores J, Velarde MC, Demaria M, Davalos AR, Wiley CD, Limbad C, et al. Cellular Senescence Promotes Skin Carcinogenesis through p38MAPK and p44/42MAPK Signaling. Cancer Res. 2020; 80(17):3606–3619.

19. Farr JN, Xu M, Weivoda MM, Monroe DG, Fraser DG, Onken JL, Negley BA, Sfeir JG, Ogrodnik MB, Hachfeld CM, LeBrasseur NK, Drake MT, Pignolo RJ, et al. Targeting cellular senescence prevents age-related bone loss in mice. Nat Med. 2017; 23(9):1072–1079.

20. Chandra A, Lagnado AB, Farr JN, Monroe DG, Park S, Hachfeld C, Tchkonia T, Kirkland JL, Khosla S, Passos JF and Pignolo RJ. Targeted Reduction of Senescent Cell Burden Alleviates Focal Radiotherapy-Related Bone Loss. J Bone Miner Res. 2020; 35(6):1119–1131.

21. Yao Z, Murali B, Ren Q, Luo X, Faget DV, Cole T, Ricci B, Thotala D, Monahan J, van Deursen JM, Baker D, Faccio R, Schwarz JK, et al. Therapy-Induced Senescence Drives Bone Loss. Cancer Res. 2020; 80(5):1171–1182.

22. Reid IR, Green JR, Lyles KW, Reid DM, Trechsel U, Hosking DJ, Black DM, Cummings SR, Russell RGG and Eriksen EF. Zoledronate. Bone. 2020; 137:115390.

23. Black DM, Delmas PD, Eastell R, Reid IR, Boonen S, Cauley JA, Cosman F, Lakatos P, Leung PC, Man Z, Mautalen C, Mesenbrink P, Hu H, et al. Once-yearly zoledronic acid for treatment of postmenopausal osteoporosis. N Engl J Med. 2007; 356(18):1809–1822.

24. Lyles KW, Colón-Emeric CS, Magaziner JS, Adachi JD, Pieper CF, Mautalen C, Hyldstrup L, Recknor C, Nordsletten L, Moore KA, Lavecchia C, Zhang J, Mesenbrink P, et al. Zoledronic acid and clinical fractures and mortality after hip fracture. N Engl J Med. 2007; 357(18):1799–1809.

25. Cengiz Ö, Polat G, Karademir G, Tunç OD, Erdil M, Tuncay I and Sen C. Effects of Zoledronate on Mortality and Morbidity after Surgical Treatment of Hip Fractures. Adv Orthop. 2016; 2016:3703482.

26. Reid IR, Horne AM, Mihov B, Stewart A, Garratt E, Wong S, Wiessing KR, Bolland MJ, Bastin S and Gamble GD. Fracture Prevention with Zoledronate in Older Women with Osteopenia. N Engl J Med. 2018; 379(25):2407–2416.

27. Reid IR, Horne AM, Mihov B, Stewart A, Garratt E, Bastin S and Gamble GD. Effects of Zoledronate on Cancer, Cardiac Events, and Mortality in Osteopenic Older Women. J Bone Miner Res. 2020; 35(1):20–27.

28. Wolfe F, Bolster MB, O’Connor CM, Michaud K, Lyles KW and Colón-Emeric CS. Bisphosphonate use is associated with reduced risk of myocardial infarction in patients with rheumatoid arthritis. J Bone Miner Res. 2013; 28(5):984–991.

29. Kranenburg G, Bartstra JW, Weijmans M, de Jong PA, Mali WP, Verhaar HJ, Visseren FLJ and Spiering W. Bisphosphonates for cardiovascular risk reduction: A systematic review and meta-analysis. Atherosclerosis. 2016; 252:106–115.

30. Cummings SR, Lui LY, Eastell R and Allen IE. Association Between Drug Treatments for Patients With Osteoporosis and Overall Mortality Rates: A Meta-analysis. JAMA Intern Med. 2019; 179(11):1491–1500.

31. Varela I, Pereira S, Ugalde AP, Navarro CL, Suárez MF, Cau P, Cadiñanos J, Osorio FG, Foray N, Cobo J, de Carlos F, Lévy N, Freije JM, et al. Combined treatment with statins and aminobisphosphonates extends longevity in a mouse model of human premature aging. Nat Med. 2008; 14(7):767–772.

32. Jeong J, Lee KS, Choi YK, Oh YJ and Lee HD. Preventive effects of zoledronic acid on bone metastasis in mice injected with human breast cancer cells. J Korean Med Sci. 2011; 26(12):1569–1575.

33. Ubellacker JM, Haider MT, DeCristo MJ, Allocca G, Brown NJ, Silver DP, Holen I and McAllister SS. Zoledronic acid alters hematopoiesis and generates breast tumor-suppressive bone marrow cells. Breast Cancer Res. 2017; 19(1):23.

34. Misra J, Mohanty ST, Madan S, Fernandes JA, Hal Ebetino F, Russell RG and Bellantuono I. Zoledronate Attenuates Accumulation of DNA Damage in Mesenchymal Stem Cells and Protects Their Function. Stem Cells. 2016; 34(3):756–767.

35. Hain BA, Jude B, Xu H, Smuin DM, Fox EJ, Elfar JC and Waning DL. Zoledronic Acid Improves Muscle Function in Healthy Mice Treated with Chemotherapy. J Bone Miner Res. 2020; 35(2):368–381.

36. Essex AL, Pin F, Huot JR, Bonewald LF, Plotkin LI and Bonetto A. Bisphosphonate Treatment Ameliorates Chemotherapy-Induced Bone and Muscle Abnormalities in Young Mice. Front Endocrinol (Lausanne). 2019; 10:809.

37. Fuhrmann-Stroissnigg H, Ling YY, Zhao J, McGowan SJ, Zhu Y, Brooks RW, Grassi D, Gregg SQ, Stripay JL, Dorronsoro A, Corbo L, Tang P, Bukata C, et al. Identification of HSP90 inhibitors as a novel class of senolytics. Nat Commun. 2017; 8(1):422.

38. Fuhrmann-Stroissnigg H, Santiago FE, Grassi D, Ling Y, Niedernhofer LJ and Robbins PD. SA-β-Galactosidase-Based Screening Assay for the Identification of Senotherapeutic Drugs. J Vis Exp. 2019; (148).

39. Li P, Zhao Z, Wang L, Jin X, Shen Y, Nan C and Liu H. Minimally effective concentration of zoledronic acid to suppress osteoclasts in vitro. Exp Ther Med. 2018; 15(6):5330–5336.

40. Indrayanto G, Putra GS and Suhud F. (2021). Chapter Six - Validation of in-vitro bioassay methods: Application in herbal drug research. In: Al-Majed AA, ed. Profiles of Drug Substances, Excipients and Related Methodology: Academic Press), pp. 273–307.

41. Gregg SQ, Robinson AR and Niedernhofer LJ. Physiological consequences of defects in ERCC1-XPF DNA repair endonuclease. DNA Repair (Amst). 2011; 10(7):781–791.

42. Farr JN, Fraser DG, Wang H, Jaehn K, Ogrodnik MB, Weivoda MM, Drake MT, Tchkonia T, LeBrasseur NK, Kirkland JL, Bonewald LF, Pignolo RJ, Monroe DG, et al. Identification of Senescent Cells in the Bone Microenvironment. J Bone Miner Res. 2016; 31(11):1920–1929.

43. Schafer MJ, Zhang X, Kumar A, Atkinson EJ, Zhu Y, Jachim S, Mazula DL, Brown AK, Berning M, Aversa Z, Kotajarvi B, Bruce CJ, Greason KL, et al. The senescence-associated secretome as an indicator of age and medical risk. JCI Insight. 2020; 5(12).

44. de Sousa FRN, de Sousa Ferreira VC, da Silva Martins C, Dantas HV, de Sousa FB, Girão-Carmona VCC, Goes P, de Castro Brito GA and de Carvalho Leitão RF. The effect of high concentration of zoledronic acid on tooth induced movement and its repercussion on root, periodontal ligament and alveolar bone tissues in rats. Sci Rep. 2021; 11(1):7672.

45. Cheng YT, Liao J, Zhou Q, Huo H, Zellmer L, Tang ZL, Ma H, Hong W and Liao DJ. Zoledronic acid modulates osteoclast apoptosis through activation of the NF-κB signaling pathway in ovariectomized rats. Exp Biol Med (Maywood). 2021; 246(15):1727–1739.

46. Khosla S. Odanacatib: location and timing are everything. J Bone Miner Res. 2012; 27(3):506–508.

47. Hewitt G, Jurk D, Marques FD, Correia-Melo C, Hardy T, Gackowska A, Anderson R, Taschuk M, Mann J and Passos JF. Telomeres are favoured targets of a persistent DNA damage response in ageing and stress-induced senescence. Nat Commun. 2012; 3:708.

48. Ubellacker JM, Baryawno N, Severe N, DeCristo MJ, Sceneay J, Hutchinson JN, Haider MT, Rhee CS, Qin Y, Gregory WM, Garrido-Castro AC, Holen I, Brown JE, et al. Modulating Bone Marrow Hematopoietic Lineage Potential to Prevent Bone Metastasis in Breast Cancer. Cancer Res. 2018; 78(18):5300–5314.

49. Takamatsu Y, Simmons PJ, Moore RJ, Morris HA, To LB and Lévesque JP. Osteoclast-mediated bone resorption is stimulated during short-term administration of granulocyte colony-stimulating factor but is not responsible for hematopoietic progenitor cell mobilization. Blood. 1998; 92(9):3465–3473.

50. Li S, Zhai Q, Zou D, Meng H, Xie Z, Li C, Wang Y, Qi J, Cheng T and Qiu L. A pivotal role of bone remodeling in granulocyte colony stimulating factor induced hematopoietic stem/progenitor cells mobilization. J Cell Physiol. 2013; 228(5):1002–1009.

51. Jacome-Galarza CE, Lee SK, Lorenzo JA and Aguila HL. Identification, characterization, and isolation of a common progenitor for osteoclasts, macrophages, and dendritic cells from murine bone marrow and periphery. J Bone Miner Res. 2013; 28(5):1203–1213.

52. Jacquin C, Gran DE, Lee SK, Lorenzo JA and Aguila HL. Identification of multiple osteoclast precursor populations in murine bone marrow. J Bone Miner Res. 2006; 21(1):67–77.

53. Saul D, Kosinsky RL, Atkinson EJ, Doolittle ML, Zhang X, LeBrasseur NK, Pignolo RJ, Robbins PD, Niedernhofer LJ, Ikeno Y, Jurk D, Passos JF, Hickson LJ, et al. A new gene set identifies senescent cells and predicts senescence-associated pathways across tissues. Nat Commun. 2022; 13(1):4827.

54. Fridman AL and Tainsky MA. Critical pathways in cellular senescence and immortalization revealed by gene expression profiling. Oncogene. 2008; 27(46):5975–5987.

55. Doolittle ML, Saul D, Kaur J, Rowsey JL, Vos SJ, Pavelko KD, Farr JN, Monroe DG and Khosla S. Multiparametric senescent cell phenotyping reveals CD24 osteolineage cells as targets of senolytic therapy in the aged murine skeleton. bioRxiv. 2023.

56. Van Gassen S, Callebaut B, Van Helden MJ, Lambrecht BN, Demeester P, Dhaene T and Saeys Y. FlowSOM: Using self-organizing maps for visualization and interpretation of cytometry data. Cytometry A. 2015; 87(7):636–645.

57. So S, Davis AJ and Chen DJ. Autophosphorylation at serine 1981 stabilizes ATM at DNA damage sites. J Cell Biol. 2009; 187(7):977–990.

58. Ambrosi TH, Marecic O, McArdle A, Sinha R, Gulati GS, Tong X, Wang Y, Steininger HM, Hoover MY, Koepke LS, Murphy MP, Sokol J, Seo EY, et al. Aged skeletal stem cells generate an inflammatory degenerative niche. Nature. 2021; 597(7875):256–262.

59. Su W, Liu G, Mohajer B, Wang J, Shen A, Zhang W, Liu B, Guermazi A, Gao P, Cao X, Demehri S and Wan M. Senescent preosteoclast secretome promotes metabolic syndrome associated osteoarthritis through cyclooxygenase 2. Elife. 2022; 11.

60. Santhanam L, Liu G, Jandu S, Su W, Wodu BP, Savage W, Poe A, Liu X, Alexander LM, Cao X and Wan M. Skeleton-secreted PDGF-BB mediates arterial stiffening. J Clin Invest. 2021; 131(20).

61. Hayflick L and Moorhead PS. The serial cultivation of human diploid cell strains. Exp Cell Res. 1961; 25:585–621.

62. Chen Z, Cordero J, Alqarni AM, Slack C, Zeidler MP and Bellantuono I. Zoledronate extends healthspan and survival via the mevalonate pathway in a FOXO-dependent manner. J Gerontol A Biol Sci Med Sci. 2021.

63. Lawson MA, Paton-Hough JM, Evans HR, Walker RE, Harris W, Ratnabalan D, Snowden JA and Chantry AD. NOD/SCID-GAMMA mice are an ideal strain to assess the efficacy of therapeutic agents used in the treatment of myeloma bone disease. PLoS One. 2015; 10(3):e0119546.

64. Rogers MJ, Mönkkönen J and Munoz MA. Molecular mechanisms of action of bisphosphonates and new insights into their effects outside the skeleton. Bone. 2020; 139:115493.

65. Clynes MA, Harvey NC, Curtis EM, Fuggle NR, Dennison EM and Cooper C. The epidemiology of osteoporosis. Br Med Bull. 2020; 133(1):105–117.

66. Farr JN, Rowsey JL, Eckhardt BA, Thicke BS, Fraser DG, Tchkonia T, Kirkland JL, Monroe DG and Khosla S. Independent Roles of Estrogen Deficiency and Cellular Senescence in the Pathogenesis of Osteoporosis: Evidence in Young Adult Mice and Older Humans. J Bone Miner Res. 2019; 34(8):1407–1418.

67. Kelly NH, Schimenti JC, Patrick Ross F and van der Meulen MC. A method for isolating high quality RNA from mouse cortical and cancellous bone. Bone. 2014; 68:1–5.

68. Mayeuf-Louchart A, Hardy D, Thorel Q, Roux P, Gueniot L, Briand D, Mazeraud A, Bouglé A, Shorte SL, Staels B, Chrétien F, Duez H and Danckaert A. MuscleJ: a high-content analysis method to study skeletal muscle with a new Fiji tool. Skelet Muscle. 2018; 8(1):25.

69. Eckhardt BA, Rowsey JL, Thicke BS, Fraser DG, O’Grady KL, Bondar OP, Hines JM, Singh RJ, Thoreson AR, Rakshit K, Lagnado AB, Passos JF, Vella A, et al. Accelerated osteocyte senescence and skeletal fragility in mice with type 2 diabetes. JCI Insight. 2020; 5(9).

70. Weivoda MM, Chew CK, Monroe DG, Farr JN, Atkinson EJ, Geske JR, Eckhardt B, Thicke B, Ruan M, Tweed AJ, McCready LK, Rizza RA, Matveyenko A, et al. Identification of osteoclast-osteoblast coupling factors in humans reveals links between bone and energy metabolism. Nat Commun. 2020; 11(1):87.

71. Subramanian A, Tamayo P, Mootha VK, Mukherjee S, Ebert BL, Gillette MA, Paulovich A, Pomeroy SL, Golub TR, Lander ES and Mesirov JP. Gene set enrichment analysis: a knowledge-based approach for interpreting genome-wide expression profiles. Proc Natl Acad Sci U S A. 2005; 102(43):15545–15550.

72. Mootha VK, Lindgren CM, Eriksson KF, Subramanian A, Sihag S, Lehar J, Puigserver P, Carlsson E, Ridderstråle M, Laurila E, Houstis N, Daly MJ, Patterson N, et al. PGC-1alpha-responsive genes involved in oxidative phosphorylation are coordinately downregulated in human diabetes. Nat Genet. 2003; 34(3):267–273.

73. Nowicka M, Krieg C, Crowell HL, Weber LM, Hartmann FJ, Guglietta S, Becher B, Levesque MP and Robinson MD. CyTOF workflow: differential discovery in high-throughput high-dimensional cytometry datasets. F1000Res. 2017; 6:748.

74. Wilson DJ. The harmonic mean p-value for combining dependent tests. Proc Natl Acad Sci U S A. 2019; 116(4):1195–1200.

