## Supplementary Figures and Table for "*In vitro* and *in vivo* effects of zoledronate on senescence and senescence-associated secretory phenotype markers"

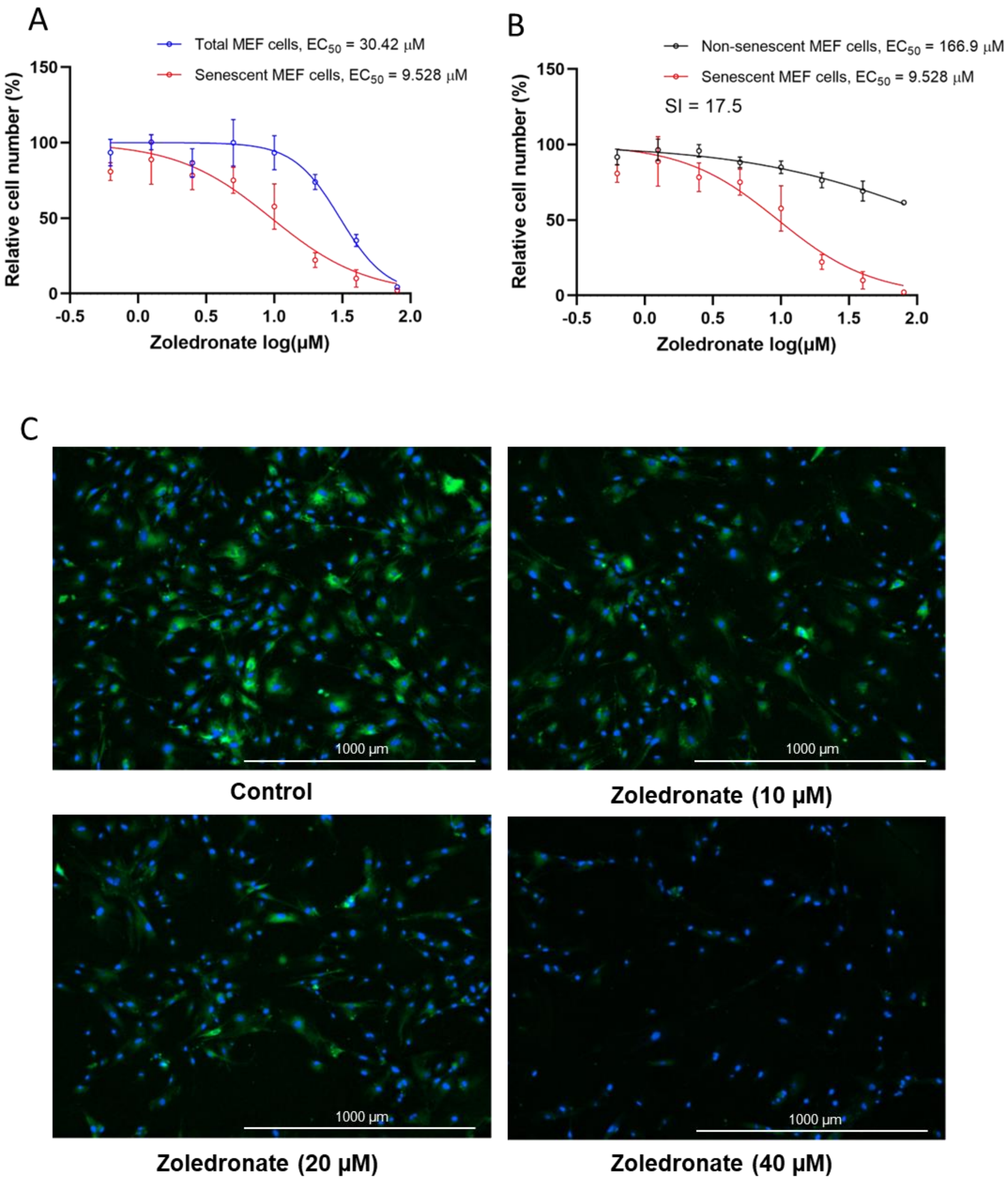

Suppl. Fig. 1. *Zoledronate has senolytic effects in mouse embryonic fibroblast (MEF) cells*. (A) Increasing concentrations (0.63-80 μM) of zoledronate were tested for 48 h in MEF cells. The figure shows the percentage of total cells (blue) and senescent *Ercc1*$^{-/-}$ MEF cells remaining following treatment. n = 3, mean ± SD; (B) Percentage of WT, non-senescent MEF cells (black) and senescent *Ercc1*$^{-/-}$ MEF cells (red) remaining after 48 hours of treatment. SI: selectivity index. n = 3; (C) Representative images of $C_{12}$FDG-based senescence assay of zoledronate in senescent *Ercc1*$^{-/-}$ MEFs. Blue fluorescence indicates nuclear staining with Hoechst 33324, and bright green fluorescence indicates SA-β-gal positive senescent cells whereas dim green fluorescence represents SA-β-gal low or negative, non-senescent cells. Images were taken using Cytation 1 at 4X.

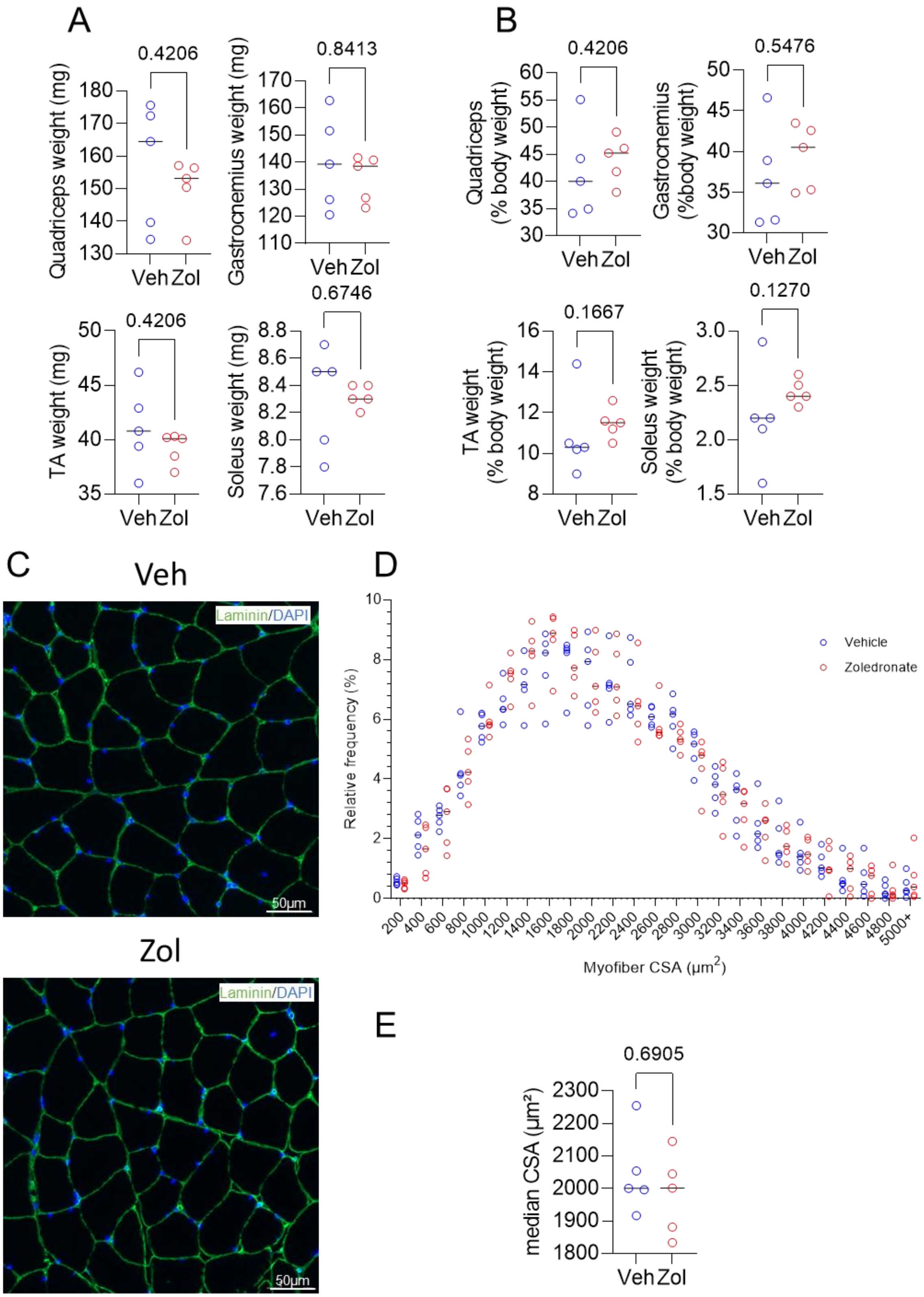

Suppl. Fig. 2. *Effects of zoledronate on muscle weight and muscle fiber cross-sectional area.* (A) Quadriceps femoris, gastrocnemius, tibialis anterior (TA), and soleus muscle weights in the vehicle- and zoledronate-treated mice; (B) Muscle weights normalized to body weights; (C) Cross-section of quadriceps muscle fibers showing laminin and nuclear staining; (D) Myofiber CSA distribution in the vehicle- and zoledronate-treated mice; (E) Quadriceps muscle CSA in the vehicle- and zoledronate-treated mice. *p*-values according to Mann-Whitney test, n=5 mice per group.

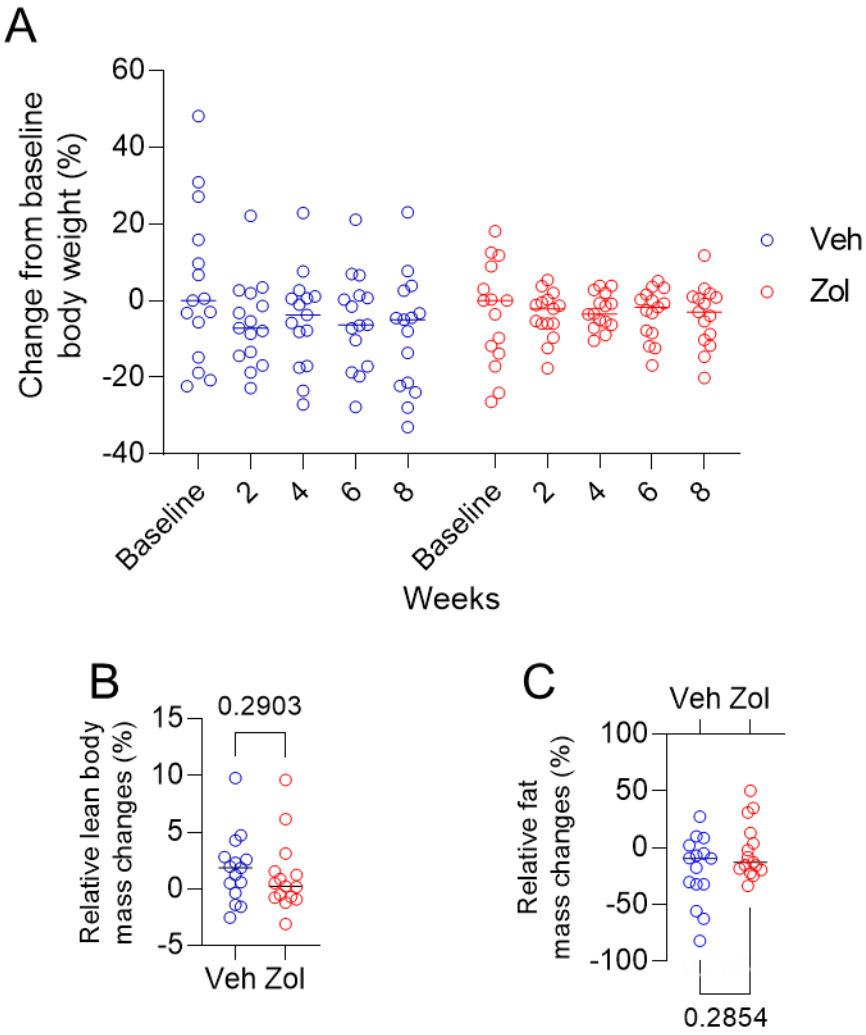

<u>Suppl. Fig. 3.</u> *Effects of Zoledronate on body composition.* (A) Baseline and weekly percent changes in body weight in the vehicle- and zoledronate-treated mice; percent change over the course of the study in (B) lean mass and (C) fat mass in the vehicle- and zoledronate-treated mice. *p*-values according to Mann-Whitney test, n=15 mice/group.

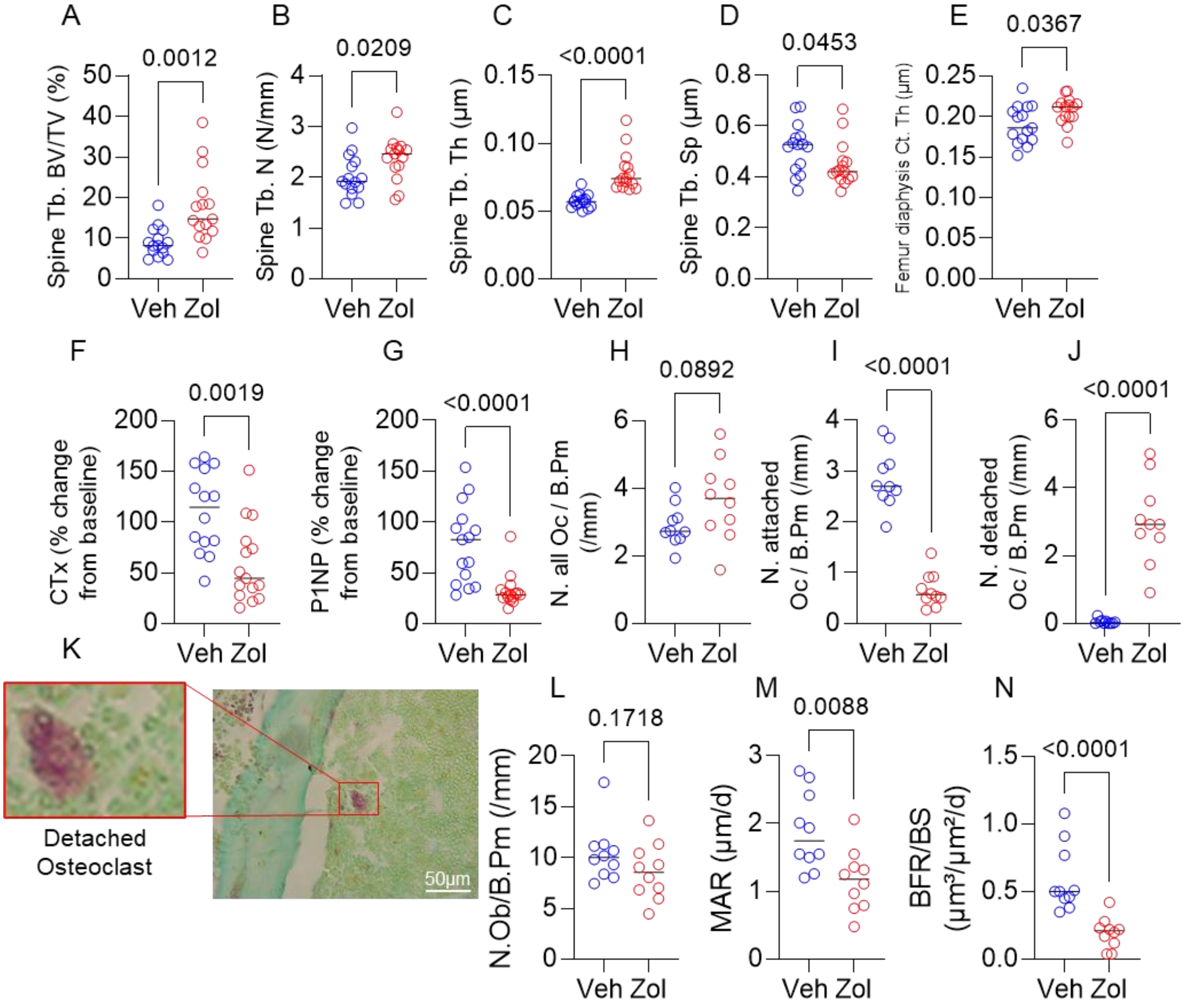

Suppl. Fig. 4. *Skeletal effects of zoledronate.* Effects of zoledronate on (A-D) spine trabecular and (E) femur diaphysis cortical parameters; (F, G), percent changes in serum CTx and PINP levels in the zoledronate- and vehicle-treated mice; (H) total osteoclasts, (I) attached osteoclasts, and (J) detached osteoclasts in the zoledronate- and vehicle-treated mice; (K) shows an example of a detached osteoclast in the zoledronate-treated mice; trabecular (L) osteoblast numbers, (M) mineral apposition rate, and (N) bone formation rate in the vehicle- and zoledronate-treated mice. *p*-values are using Mann-Whitney test; n = 10-15 mice/group.

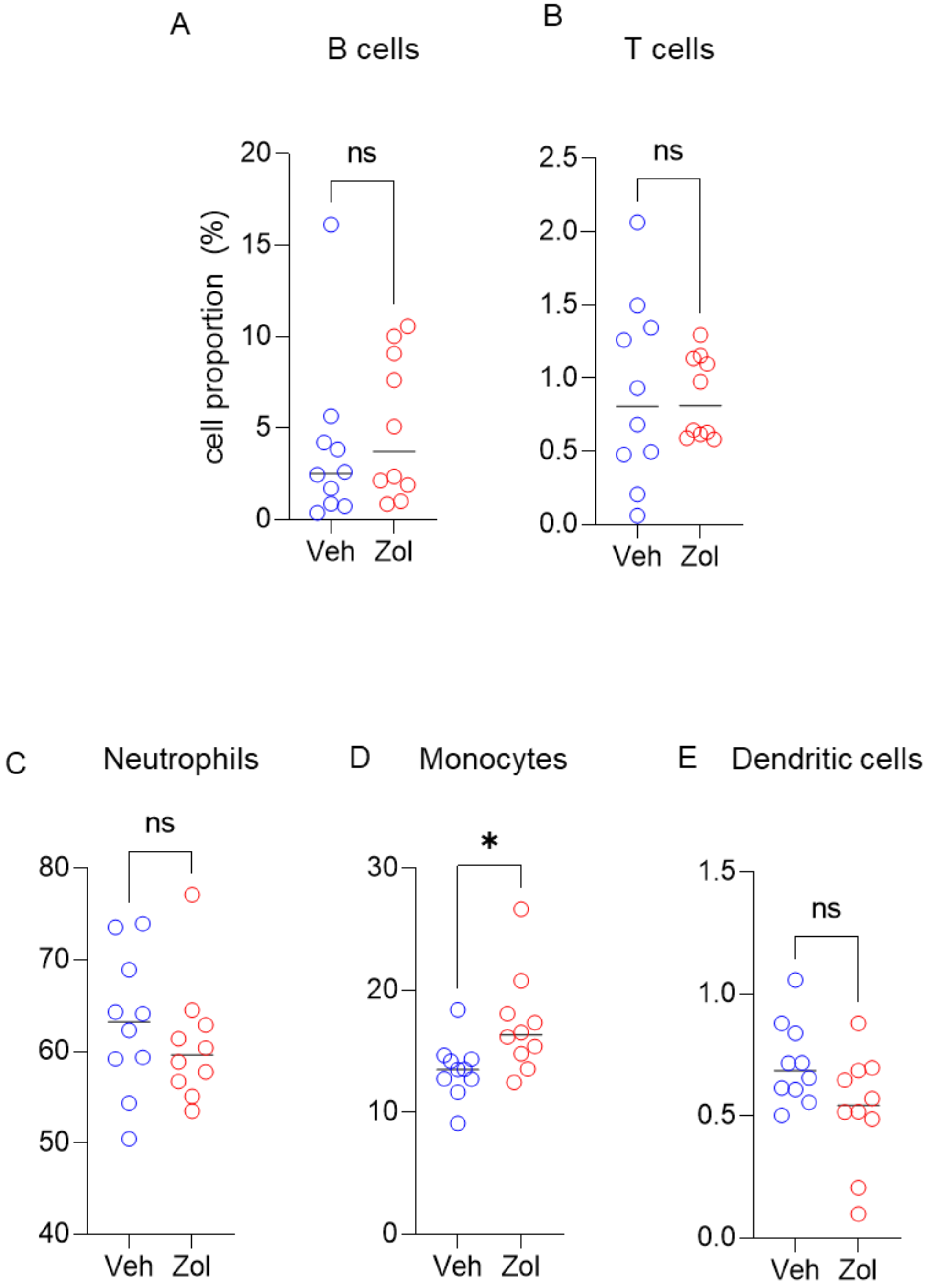

<u>Suppl. Fig. 5.</u> *Zoledronate does not reduce other bone marrow hematopoietic cell populations.* Percentages of (A) B-cells, (B) T-cells, (C) neutrophils, (D) monocytes, or (D) dendritic cells are not altered by zoledronate. N=10 mice in the control and n=10 in the zoledronate group.

Suppl. Table 1. Antibodies used for CyTOF, conjugated metals, supplier, and identification numbers.

| **Reagent** | **Conjugated Metal** | **Source** | **Identifier** |
|---|---|---|---|
| Hematopoietic | | | |
| CD45 | 089Y | Fluidigm | 3089005B |
| Stem/Progenitor | | | |
| CD34 | 163Dy | Thermo Fisher | MA5-17826 |
| Emcn | 164Dy | ThermoFisher | 14-5851-82 |
| CD117 | 173Yb | FDM | 3173004B |
| Granulocytes | | | |
| CD11b | 111Cd | Biolegend | 101201 |
| Macrophage | | | |
| F4-80 | 113Cd | Biolegend | 123101 |
| CX3CR1 | 161Dy | Biolegend | 149002 |
| Monocytes | | | |
| Ly6C | 141Pr | Biolegend | 128039 |
| CD11c | 142Nd | FDM | 3142003B |
| CCR2 | 156Gd | Novus | MAB55381-100 |
| CD115 | 174Yb | Biolegend | 135521 |
| CD14 | 165Ho | Biolegend | 123321 |
| T cells | | | |
| CD4 | 145Nd | FDM | 3145002B |
| CD3e | 152Sm | FDM | 3152004B |
| CD8a | 168Er | FDM | 3168003B |
| Osteoclasts | | | |
| Ctsk | 147Sm | abcam | ab37259 |
| ACP5 | 151Eu | Abcam | ab83050 |
| B cells | | | |
| CD45R | 153Eu | Biolegend | 103249 |
| Neutrophils | | | |
| Catalase | 158Gd | Abcam | ab223793 |
| Ly6G | 170Er | Biolegend | 127637 |

| | | | |
|---|---|---|---|
| B cells | | | |
| CD19 | 166Er | FDM | 3166015B |
| BMSCs | | | |
| Ly-6A | 169Tm | FDM | 3169015B |
| SASP | | | |
| IL-1α | 171Yb | Biolegend | 503202 |
| IL-1β | 159Tb | Cell Signaling | 31202 |
| CXCL1 | 175Lu | R&D Systems | MAB453-500 |
| pNFkB | 110Cd | Abcam | ab16502 |
| MCP-1 | 149Sm | Thermo Fisher | MA5-17040 |
| PAI-1 | 160Gd | Abcam | ab125687 |
| IL-6 | 167Er | FDM | 3167003B |
| Senescence | | | |
| p53 | 116Cd | Abcam | ab252388 |
| p16 | 155Gd | Abcam | ab232402 |
| p21 | 176Yb | Santa Cruz | sc-6246 |
| DNA Damage | | | |
| pATM | 114Cd | Invitrogen | 14-0146-82 |
